# From background to foreground: secondary antibodies coupled to lipophilic ATTO dyes enable high-density membrane labeling in super-resolution and expansion microscopy

**DOI:** 10.64898/2026.07.10.737767

**Authors:** Jim Dompierre, Silvia del Pozo Perera, Louis Hurson, Arnaud Mourier, Anne Devin, Manuel Rojo

## Abstract

Classical immunolabeling approaches can achieve homogeneous and continuous labeling of cellular membranes and organelles at wide-field and confocal resolution. In super-resolution and expansion microscopy, however, the lack of high-density labels hampers the localization of membrane proteins and protein complexes within their membrane context. Here we show that secondary antibodies coupled to the lipophilic dyes ATTO 647N or ATTO 550 brightly label the nuclear envelope, mitochondria, and endoplasmic reticulum of fixed, permeabilized cells, and that graded labelling intensities allow selective visualization of organelles and precise segmentation of mitochondria. Using state-of-the-art super-resolution and expansion microscopy, we achieve high-density labelling of nuclear and mitochondrial membranes, with targeting and density comparable to existing membrane-labelling approaches and a signal that can be further amplified with additional secondary antibodies. Finally, we show that these dye-conjugated IgG allow to resolve mitochondria-ER contacts and mitochondrial ultrastructure as well as precise visualization of the nuclear envelope and its invaginations. This study demonstrates that secondary antibodies conjugated to lipophilic fluorophores represent stable, convenient and affordable tools for organelle visualization in conventional microscopy and for high-density labeling of membranes in super-resolution and expansion microscopy.

## Introduction

In fluorescence microscopy, proteins of interest are localized by complementary strategies including the detection of endogenous proteins with specific antibodies and/or the expression of exogenous proteins carrying epitope and/or fluorescent tags. The simultaneous co-labeling of cellular compartments, organelles and/or membranes is required for precise determination of their local and subcellular distribution. Co-visualization of cellular organelles and membranes can be achieved by the use of fluorescent organic dyes accumulating to specific membranes and/or organelles in cultured cells (*e.g.* (Poot et al., 1996; Freundt et al., 2007; Fam et al., 2018; Collot et al., 2019)). However, the selectivity and/or efficacy of certain vital dyes can be limited and/or altered by the metabolic and functional status of cells and organelles. The targeting of mitochondrial dyes in cultured cells, for example, often depends on the membrane potential, leading to diminished or abolished labeling of dysfunctional organelles (Poot et al., 1996). In contrast, the labeling of organelles and membranes with antibodies depends on the abundance of marker proteins and on the availability of antibodies detecting them in fixed and permeabilized cells (Malka et al., 2007). The expression of proteins of known localization carrying epitope or fluorescent tags is also suited for labeling organelles but (over)expression can lead to protein mistargeting and/or alteration of the targeted compartment (*e.g.* (Snapp et al., 2003)).

The visualization of biological membranes and of their continuity depends on the density of the labeled or labeling molecule. At the limited resolution of wide-field and confocal microscopy, the accumulation of organic dyes and/or the localization of abundant endogenous proteins by immunofluorescence can be sufficient to display apparently continuous membranes of cells or organelles. At the significantly higher resolution achieved by super-resolution techniques such as Structured Illumination Microscopy (SIM), Stimulated Emission Depletion microscopy (STED), Stochastic Optical Reconstruction Microscopy (STORM), or expansion microscopy (ExM), however, the sparse distribution of fluorescent dyes, proteins and protein complexes hampers the visualization of membrane continuity (Lambert and Waters, 2017). Although imaging algorithms can generate faithful reconstructions of cellular and organellar membranes (Marin et al., 2023), it would be extremely advantageous to identify dyes and tools labeling biological membranes with higher density.

A majority of the organic dyes that allow specific labelling of membranes and/or organelles enable their visualization in living cells, as well as the co-localization of fluorescently tagged proteins. However, only a minority of them withstands cellular fixation and/or membrane permeabilization, two steps required for co-localization of proteins by immunofluorescence microscopy. Conventional mitochondrial dyes of the MitoTracker^TM^ series carry a chloromethyl group for covalent linkage to proteins (Poot et al., 1996)) and a STED-compatible mitochondrial dye (PKMO FX) carries an amino-group for aldehyde fixation (Chen et al., 2024). Other membrane-dyes (mCLING, pGK13a) are based on the linkage of a membrane-anchoring palmitoyl chain to a poly-lysine peptide enabling aldehyde-fixation (Revelo et al., 2014) (Shin et al., 2025) (Zhuang et al., 2026) or on the addition of a reactive ester to membrane anchored dyes (MemGraft) (Aknine et al., 2025). A third strategy, specific for ExM, relies on the addition of acryl-moieties immobilizing dyes by direct polymerization into the ExM-gel (Wen et al., 2020).

In this work, we describe a strategy for high-density labeling of cellular membranes in fixed cells with common, stable and readily available reagents, namely fluorescent secondary antibodies that are directly bound to membranes by coupled lipophilic dyes (ATTO 647N or ATTO 550). Widefield microscopy shows that these IgG molecules label mitochondria, endoplasmic reticulum and nuclear envelope of fixed and permeabilized cells. Super-resolution and expansion microscopy show that these IgG molecules enable high density labeling and ultrastructural characterization of mitochondrial and nuclear membranes.

## Results

### Secondary antibodies coupled to lipophilic ATTO 647N or ATTO 550 label cellular structures of fixed and permeabilized cells

During immunofluorescence experiments with established protocols (Malka et al., 2007), we made use of secondary antibodies labeled with ATTO 647N (IgG-A647N), a lipophilic dye known for its high fluorescence intensity and excellent photostability. We observed an unspecific background that was negligible upon detection of abundant, brightly stained antigens but interfered with the localization of low abundance or faintly labeled proteins. An unspecific and/or mitochondrial background of secondary IgG-A647N having been previously reported (Kolmakov et al., 2010) (Wurm et al., 2010), we setup to investigate whether this background signal could be turned into a membrane-labeling strategy. Mouse Embryo Fibroblasts (MEFs) were fixed with aldehydes and permeabilized with detergents. For improved structural preservation of samples, we used a formaldehyde-based fixative containing glutaraldehyde (3,2% PFA, 0,1% GA) (Laporte et al., 2022) (Kim et al., 2022). Cells were then directly incubated with a mixture of two secondary antibodies: IgG-A647N and, as a control, IgG-ATTO 488 (IgG-A488). Unlike IgG-A488, IgG-A647N labeled intracellular structures resembling cellular membranes (Figure 1A). Intracellular structures were observed at the low concentrations applied for visualization of primary antibodies, but labeling intensity was significantly enhanced upon incubation of cells with threefold and tenfold antibody concentrations (Figure 1A, B). To confirm that this labeling was specific to IgG molecules coupled to ATTO 647N, fixed and permeabilized cells were incubated with secondary antibodies coupled to dyes with similar spectral properties (IgG-Cy5, IgG-AlexaFluor 647 and IgG-CF640R, Supplementary Fig. S1). Fluorescence microscopy revealed that none of them decorated the pattern observed with IgG-A647N or produced the strong signal observed with IgG-A647N (Fig. 1B, C). We established that labeling of cellular structures was observed with IgG-A647N of various specificities (anti-mouse, anti-goat and anti-rat IgG) that had been generated in different hosts (goat and rabbit) and were provided by different suppliers (Sigma-Aldrich and Rockland) and pursued their characterization.

**Figure 1.**
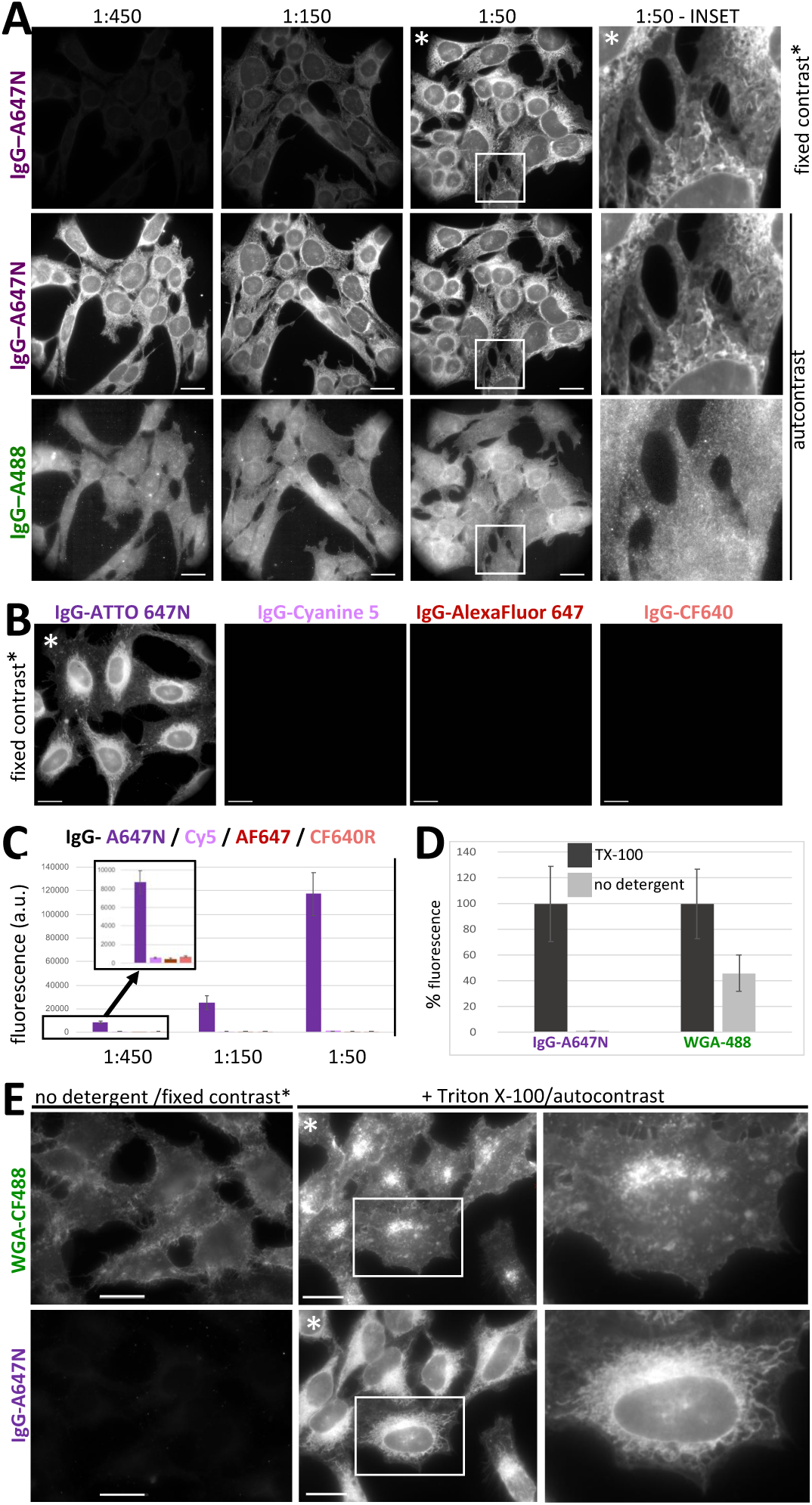
IgG-molecules coupled to ATTO 647N (IgG-A647N) label fixed and permeabilized cells. A-C: MEFs (A) or HeLa cells (B, C) were fixed with PFA/GA, permeabilized with Triton X-100 and incubated with secondary IgG coupled to the indicated fluorescent dyes at the indicated dilution (A, C) or at 1:50 dilution (B). Cells were imaged with dye-specific excitation/emission settings for identical exposure times. A, B: Images were treated with brightness/contrast settings for maximal autocontrast, or with brightness/contrast settings of the highest IgG-ATTO 647N concentration (fixed contrast*) C: Relative signal intensity in images of cells decorated with different dilutions of IgG coupled to ATTO 647N (A647N), Cyanine5 (Cy5), AlexaFluor 647 (AF647) or CF640R. Bars: 20 µm. D, E: Hela cells were fixed and subjected to detergent-mediated permeabilized (+ Triton X-100) or not (no detergent) and incubated with a mixture of IgG-A647N and WGA-CF488. Images of permeabilized and unpermeabilized cells were acquired under identical conditions. D: Quantification of fluorescence intensity in 6 independent images. E: Images of unpermeabilized cells were treated with brightness/contrast settings for permeabilized cells (fixed contrast*).

First, we investigated the effect of detergent-mediated permeabilization on direct labeling by IgG-A647N. To this end, fixed HeLa cells were incubated with a mixture of IgG-A647N and of fluorescently-labeled wheat germ agglutinin (WGA), a lectin targeting glycoproteins enriched in the medial/trans-side of the Golgi apparatus and in the plasma membrane (Tartakoff and Vassalli, 1983) (Schroeter et al., 2026). As expected, WGA-labeling was restricted to the plasma membrane in non-permeabilized cells and labeled the plasma membrane and the bright perinuclear pattern corresponding to the Golgi apparatus after detergent-mediated permeabilization (Fig. 1D, E). These results, in accordance with the known localization of WGA-targeted glycoproteins, confirmed that membrane permeabilization is dispensable for labeling of plasma-membrane glycoproteins, but required for access of lectins (molecular mass ∼40 kDa) to glycoproteins of the Golgi apparatus. Detergents modulated the labeling by IgG-A647N in a similar manner: a membrane-like pattern was labeled by IgG-A647N in permeabilized cells, but labeling was abolished, rather than reduced, in unpermeabilized cells (Fig. 1D, E). These experiments imply that labeling of cellular structures by IgG-A647N (molecular mass ∼150 kDa) requires membrane permeabilization, as in standard immunofluorescence experiments based on specific labeling with primary and secondary antibodies.

### Cellular labelling by fluorophore-conjugated IgG molecules relies on lipophilic ATTO dyes

The ATTO 647N dye being a relatively lipophilic molecule that interacts with liposome membranes (Hughes et al., 2014), we reasoned that its hydrophobicity may be responsible for direct labeling of cellular structures. To corroborate this hypothesis, we investigated the labeling capacity of IgG-molecules coupled to ATTO 550 (IgG-A550), a dye with similar molecular structure (Supplemental Fig. S1) and lipid-binding capacity (Hughes et al., 2014). We observed that, upon incubation of cells with a mixture of IgG-A647N and IgG-A550, both IgG-molecules visualized the entire cell as well as brighter cellular structures reminiscent of mitochondria and of the nuclear envelope (Fig. 2A). As observed for IgG-A647N, the labeling intensity of IgG-A550 was proportional to the antibody concentration (Fig. 2B).

**Figure 2:**
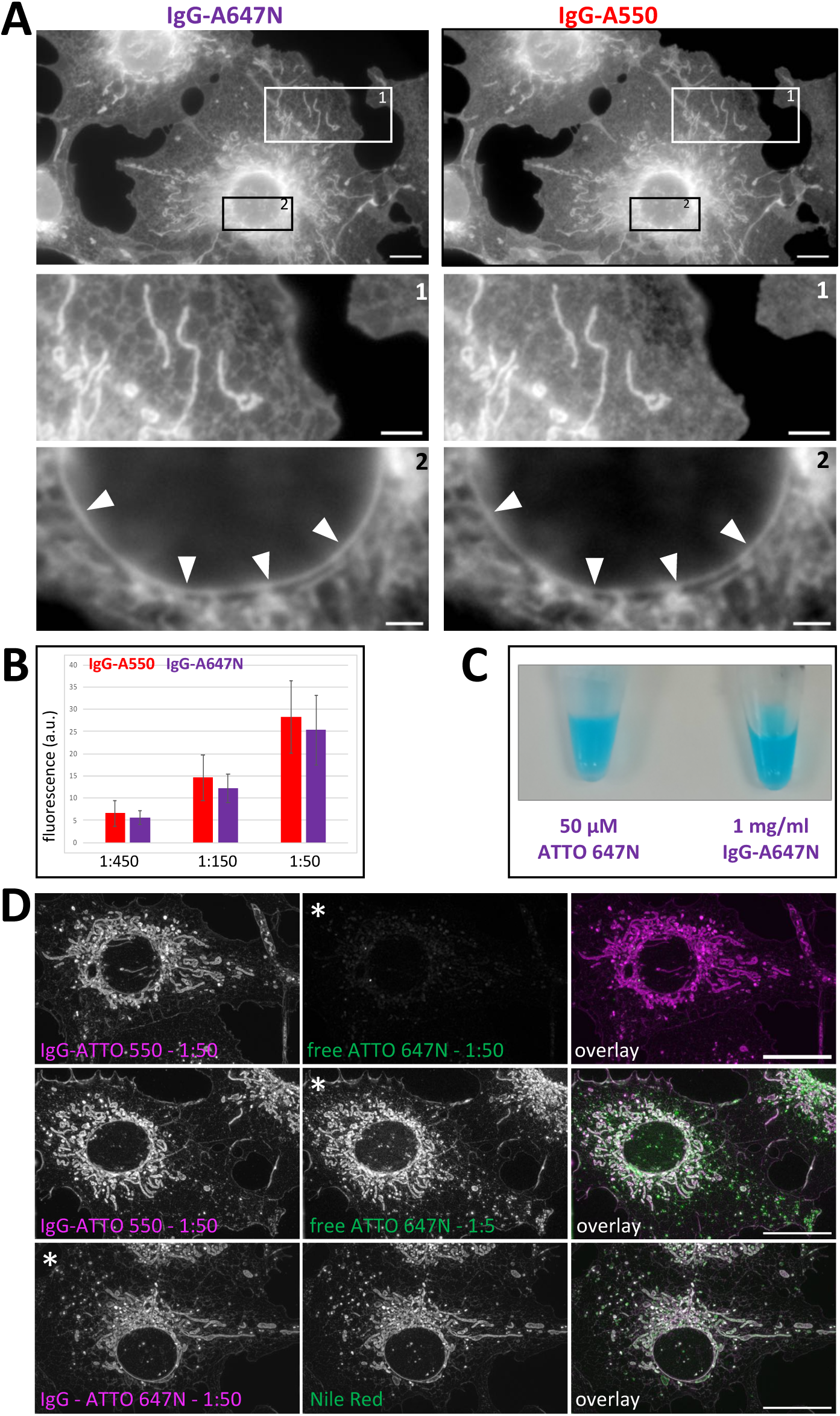
Membrane labeling by dye-coupled IgG-molecules relies on the lipophilic properties of fluorophores. A, B: COS-7 cells were fixed, permeabilized and incubated with a mixture of IgG-A647N and IgG-A550 (**A:** 1:50, **B:** indicated dilution). The position of enlarged insets 1 and 2 is indicated by white squares. The focal planes of inset 1 and 2 correspond to the optimal focal plane of the depicted structures. **B:** Relative signal intensity in images of cells decorated with increasing concentrations of IgG-A550 and IgG-A647N. **C:** Images of tubes containing free and IgG-coupled ATTO 647N at similar dye concentrations. **D:** COS-7 cells were fixed, permeabilized and incubated with the indicated dye mixtures. The images of free and IgG-cojugated ATTO 647N were acquired under identical conditions; brightness/contrast was set to the autocontrast of the images of IgG-ATTO 647N (*).

To corroborate that labeling by IgG-A647N and IgG-A550 relied on the lipophilic nature of the corresponding ATTO dyes, we compared the labeling pattern obtained with IgG-A550 with that of unconjugated ATTO 647N. Free and conjugated ATTO dyes labeled highly similar patterns (Fig. 2D), but free ATTO 647N had to be applied at tenfold higher dye concentrations to obtain labeling intensities similar to those achieved by IgG-A647N (Fig. 2C, D). Next, we compared the labeling pattern of IgG-A647N with that of Nile Red, a hydrophobic dye mainly used for the labelling of lipid droplets (Fam et al., 2018). The highly similar patterns obtained with IgG-A647N and Nile Red (Fig. 2D) confirmed that dye-hydrophobicity plays a major role in membrane labeling and suggest that fluorescently-labelled IgG molecules bind to cellular structures *via* lipophilic ATTO dyes. We further observed that, in contrast to IgG-coupled dyes, labeling by free dyes declined significantly upon transfer of labeled cells from PBS to a microscopy mounting medium. The inability to preserve labeling in microscopy mounting medium prompted us to abandon the characterization of unconjugated dyes and to pursue the study of dye-coupled IgG-molecules and their possible use as membrane labeling reagents.

### Antibodies coupled to lipophilic ATTO dyes label mitochondria, the nuclear envelope and the endoplasmic reticulum

The resemblance of the labeled structures to cellular organelles (Figs. 1, 2) prompted us to investigate their identity. Incubation of cells stably expressing GFP molecules anchored to the mitochondrial outer membrane (Fig. 3A) or of cells that had been labeled with Mitotracker Red (Supplementary Figure S2A) revealed that the structures most brightly labeled with IgG-A647N correspond to mitochondria. However, in addition to mitochondria, IgG-A647N weakly labeled reticular structures reminiscent of the ER (Fig. 3A, Supplementary Figure S2A). To establish the identity of the latter, cells labeled with IgG-A647N were co-labeled with antibodies against calnexin, an established ER-marker, and with Concanavalin A (ConA), a lectin binding to core-glycosylated proteins that are enriched in (but not restricted to) the ER and the cis-side of the Golgi apparatus (Tartakoff and Vassalli, 1983; Schroeter et al., 2026). As expected, calnexin and ConA displayed similar distributions within the ER and the nuclear envelope (Fig. 3B). The distribution of IgG-A647N overlapped with that of both ER markers, with a stronger labeling of the NE than the ER (Fig. 3B).

**Figure 3:**
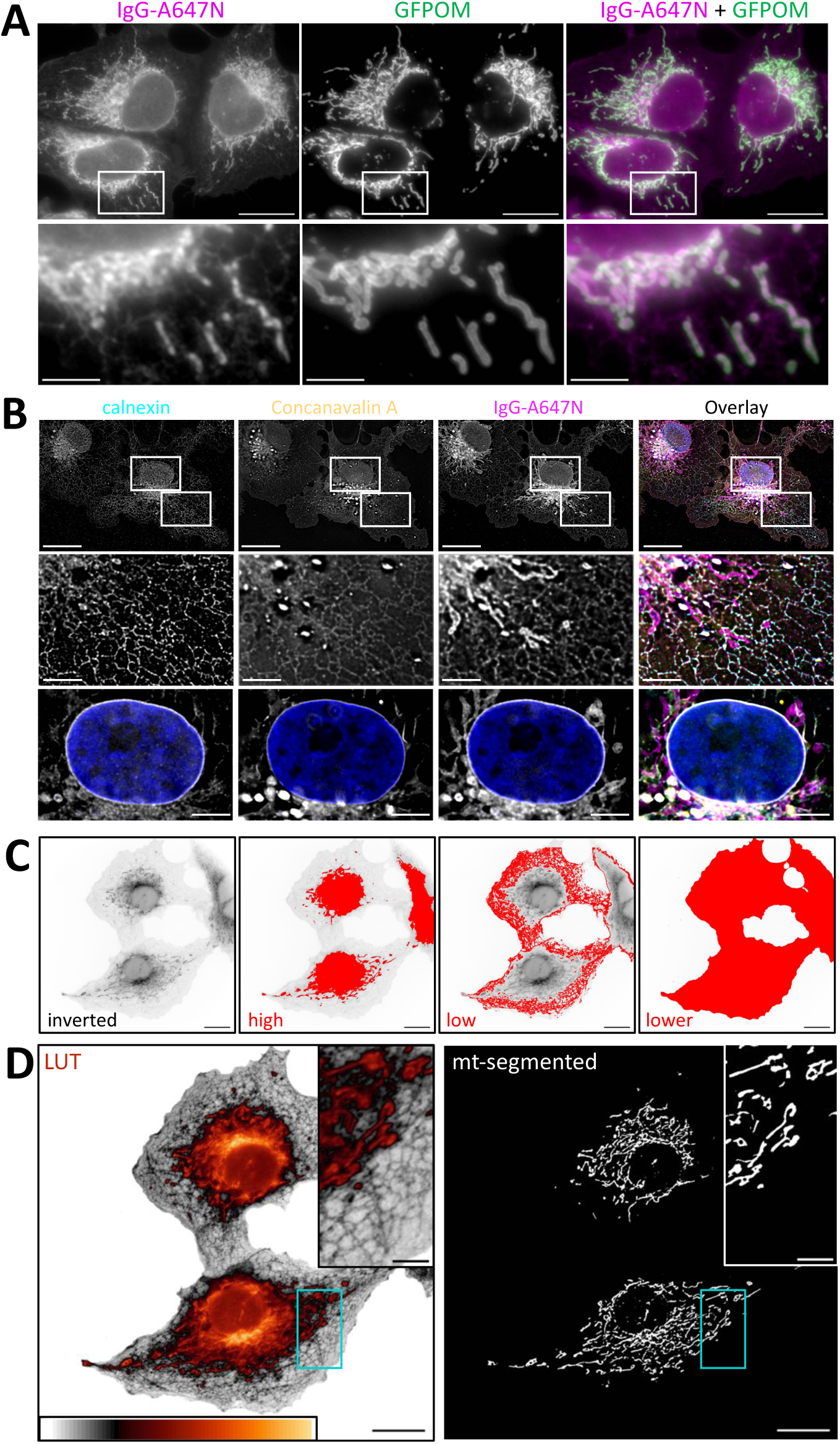
IgG-A647N labels mitochondria, nuclear envelope and endoplasmic reticulum (ER) of fixed permeabilized cells. **A**: Stably transfected 143B cells expressing EGFP targeted to the outer membrane (GFPOM) were fixed, permeabilized and decorated with IgG-A647N. The structures most brightly labeled with IgG-A647N are positive for GFPOM. **B:** COS-7 cells were fixed, permeabilized and labeled with IgG-A647N, Concanavalin-A AlexaFluor 488 and antibodies against ER-marker calnexin. Images show single planes of deconvoluted Z-stacks. **C, D:** COS-7 cells incubated with IgG-A647N after fixation and permeabilization. **C:** Inverted images treated with different levels of image thresholding. **D:** Image pseudo-colored with MQ div-autumn LUT highlights the different label intensity of mitochondria and ER. Binary image of segmented mitochondria (mt-segmented). Bars: 20 µm (main figures) and 5 µm (insets).

The ability of IgG-A647N to stain several organelles is convenient for visualization of overall cell organization. However, as it is advantageous to dispose of tools enabling the identification and segmentation of precise subcellular structures and/or organelles, we setup to investigate the capacity of IgG-A647N to image selected organelles, notably strongly labeled mitochondria. Image inversion and quantitative thresholding confirmed the differential labelling intensity of mitochondria and ER and the weaker labelling of entire cells (Fig. 3C). Next, we applied a lookup table (LUT) highlighting the differential labeling intensity of mitochondria and ER (Fig. 3D: LUT). Because mitochondria and ER labelled with IgG-A647N were distinguishable using the selected LUT, we applied Weka trainable segmentation (Arganda-Carreras et al., 2017) with default settings. This yielded precise segmentation of mitochondria, even in the crowded perinuclear region, where strong signals prevent threshold-based segmentation (Fig. 3D).

It is important to state (i) that specific labeling and segmentation represent major limiting steps for the analysis of mitochondrial distribution and morphology (Hemel et al., 2021) (Lefebvre et al., 2025), hallmarks of mitochondrial (dys)function (Guillery et al., 2008a; Barsa et al., 2025) and (ii) that the ability to visualize mitochondria after fixation, with dyes that do not depend on the mitochondrial membrane potential, is highly advantageous for mitochondrial research. Indeed, the mitochondrial accumulation of numerous vital dyes (including MitoTracker^TM^) is diminished or even abolished upon mitochondrial depolarization and/or dysfunction (Poot et al., 1996). In supplementary Figure S2B, we show that IgG-A647N enables mitochondrial imaging in cells subjected to treatments with (i) cccp, a protonophore that dissipates the mitochondrial membrane potential, inhibits inner membrane fusion and induces their fragmentation (Legros et al., 2002) (Guillery et al., 2008b) (Guillery et al., 2008a), (ii) valinomycin, a K+-specific ionophore that depolarizes mitochondria and induces their swelling and perinuclear coalescence (Malka et al., 2005; Sauvanet et al., 2012), (iii) cycloheximide, an inhibitor of cytosolic translation that drifts the fusion/fission equilibrium towards mitochondrial elongation (Tondera et al., 2009) and (iv) with 2-Deoxy-D-Glucose, an inhibitor of glycolysis that shifts energy metabolism towards oxidative phosphorylation (Guillery et al., 2008a). In conclusion, these results show that IgG-A647N is a convenient tool for fluorescent labelling of the NE, mitochondria, the ER and entire cells and that graded labelling intensities allow selective visualization of the NE and the ER and precise segmentation of mitochondria.

### Antibodies coupled to lipophilic ATTO dyes enable high density labeling of membranes in super-resolution microscopy

Having established the capacity of IgG-coupled ATTO dyes to label cells and organelles, we setup to investigate whether and how these dyes can be applied in super-resolution microscopy. Cells labeled with IgG-A647N, ConA and DAPI were first analyzed with a Machine Intelligent-Structured Illumination Microscope (MI-SIM), a SIM setup that can reach resolutions down to 60 nm (Mezache and Leterrier, 2025). The overall patterns resembled those observed by wide-field microscopy: IgG-647N strongly labeled mitochondria and NE and displayed a much weaker ER-labeling (Fig. 4A). Noteworthy, the mitochondrial labeling by IgG-A647N displayed a dense internal pattern resembling that of inner cristae membranes (Fig. 4A, arrows). The ConA lectin labeled the NE and the ER with similar intensity and, in combination with IgG-A647N, visualized mitochondria-ER contacts (Fig. 4A, arrowheads).

**Figure 4:**
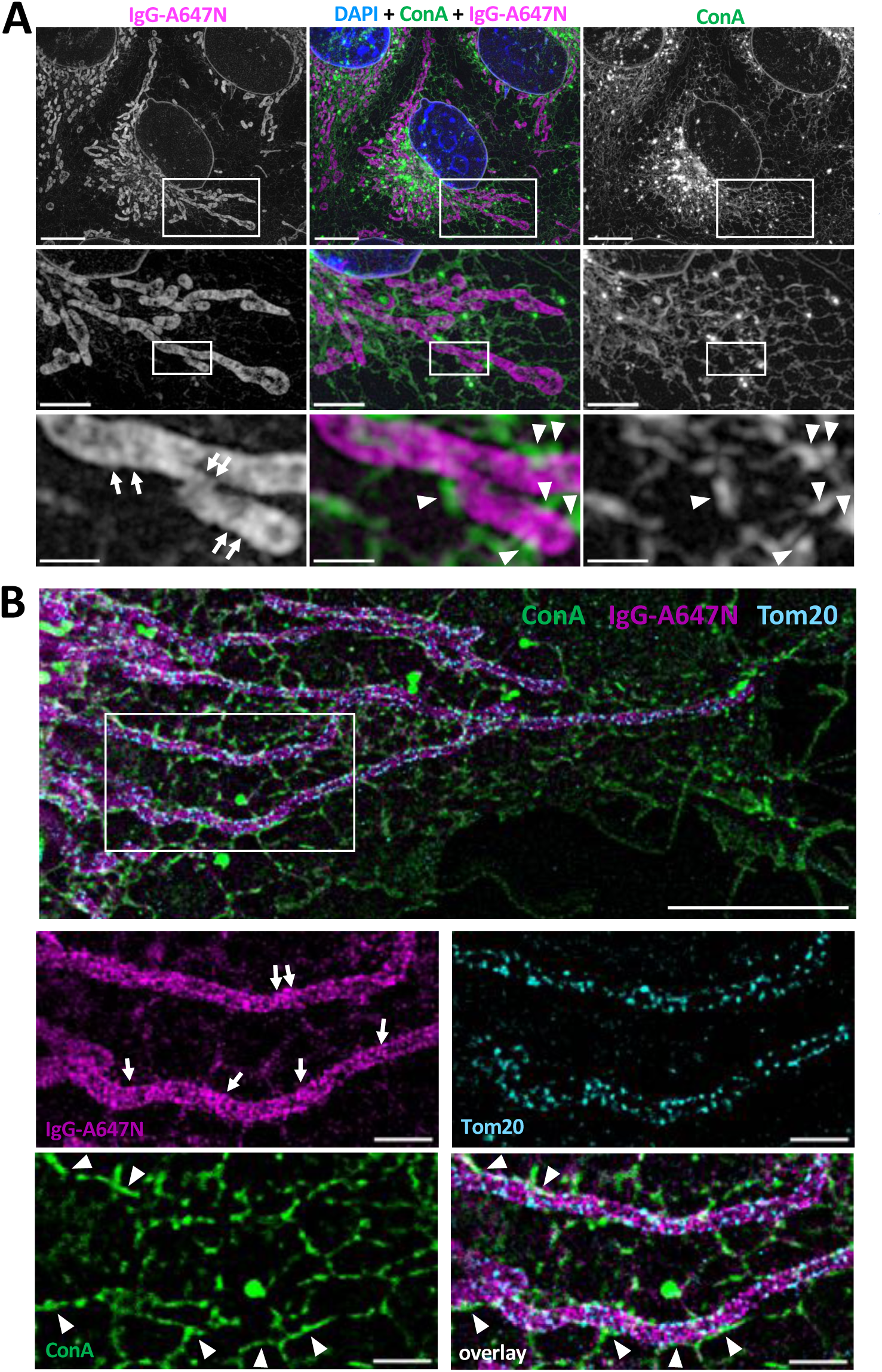
IgG-A647N enables visualization of densely labeled mitochondria by MI-SIM and STED microscopy. **A:** COS-7 cells were fixed, permeabilized and labeled with IgG-A647N, Concanavalin-A AlexaFluor 488 (ConA) and DAPI. Labeled cells were imaged by MI-SIM microscopy. Bars 10 µm (main), 4 µm (1^st^ inset), 1 µm (2^nd^ inset). **B:** HeLa cells were fixed, permeabilized and decorated with IgG-A647N, Concanavalin AlexaFluor 488 (ConA) and rabbit anti-Tom20 (Tom20). Bars 5 µm (main figure) and 1 µm (insets). Arrows point to cristae-like structures; arrowheads to mitochondria-ER contacts.

The ATTO 647N dye being well-suited for STED, a technique achieving resolutions down to 40 nm (Kolmakov et al., 2010; Wurm et al., 2010), we investigated the capacity of IgG-A647N to visualize membranes with this technique. Cells were labelled with IgG-A647N and ConA, as above, and with antibodies specific for Tom20, a component of the TOM (translocase of the outer membrane) complex. Similarly to other TOM-components localized with STED microscopy (Stoldt et al., 2019), Tom20 appeared enriched as multiple dots at the mitochondrial periphery (Fig. 4B). In contrast, IgG-A647N displayed a granular distribution within and throughout mitochondria (Fig. 4B). The distribution of the IgG-A647N dye was less homogeneous than with SIM and appeared occasionally enriched within striated structures reminiscent of inner mitochondrial cristae (Fig. 4B, arrows). As with SIM, the combination of IgG-A647N and ConA allowed to visualize mitochondria-ER contacts (Fig. 4B, arrowheads).

The suboptimal blinking properties of ATTO 550 and ATTO 647N for STORM microscopy (Dempsey et al., 2011), led us to investigate the possibility to visualize membranes ‘primarily’ labeled with IgG-A550 with ‘secondary’ antibodies carrying AlexaFluor 647, a thoroughly characterized dye perfectly suited for this microscopy approach (Dempsey et al., 2011). Permeabilized and fixed cells were sequentially incubated with goat IgG-A550 (for primary membrane labeling) and with a secondary donkey anti goat IgG-AlexaFluor 647. Analysis by conventional wide-field microscopy revealed strongly overlapping distributions of primary and secondary labels (Supplementary Fig. S3A). Imaging of IgG-AlexaFluor 647 by STORM confirmed intense mitochondrial labeling and revealed an intramitochondrial pattern reminiscent of inner membrane cristae (Supplementary Fig. S3B, C, arrows). Altogether, this series of experiments demonstrates that IgG-A647N and IgG-A550 represent potent tools for the visualization of cellular membranes and organelles, notably mitochondria, with several super-resolution microscopy techniques.

### Antibodies coupled to lipophilic ATTO dyes enable high density labeling of mitochondrial and nuclear membranes in expansion microscopy (ExM)

Next, we investigated whether membranes labeled with IgG molecules coupled to lipophilic ATTO dyes can be visualized by expansion microscopy (ExM), a super-resolution approach based on the immobilization of samples in a swellable hydrogel, their homogenization by proteases or detergents and their isotropic expansion (4X – 10X) in three dimensions (Passmore et al., 2026). We reasoned that membrane-bound IgG-A647N could be anchored to the ExM-gel (as primary and secondary antibodies recognizing specific proteins) and allow membrane-visualization after hydrogel expansion. We used a modified U-ExM recipe (Gambarotto et al., 2021) with lower bis-acrylamide content that achieved 5,6X expansion in a single step (see Materials and Methods). To investigate whether membrane-bound IgG-A647N was efficiently retained during polymerization and homogenization of ExM gels, we compared its behavior with that of Membrane ExM 561 (MemExM), a fluorescent membrane probe that carries a polymerizable acrylate group for covalent and quantitative anchoring within ExM gels (Wen et al., 2020). Cells were fixed, permeabilized and incubated with a mixture of IgG-A647N and MemExM. After polymerization, denaturation and expansion (∼5,6X), the U-ExM gel was visualized with a 40X objective (total enlargement ∼224X). Imaging revealed that the patterns labeled by IgG-A647N and MemExM were highly similar and included the NE, mitochondria, the ER and the plasma membrane (Fig. 5A). Next, we compared IgG-A647N labeling with an overall protein-staining procedure that is commonly used to visualize cellular and organellar ultrastructure, notably upon iterative expansion (M’Saad and Bewersdorf, 2020) (Louvel et al., 2023). Cells were fixed, permeabilized, labeled with IgG-A647N and subjected to polymerization into an U-ExM gel. After homogenization with SDS, gels were incubated with NHS-ATTO 488 (NHS-A488) and both dyes were visualized with a 40X objective after expansion in water (total enlargement ∼224X). Analysis by wide-field microscopy shows that IgG-A647N and NHS-A488 display similar, but not identical, labeling patterns (Fig. 5B). Both dyes labeled mitochondria and the NE (albeit with different relative intensities), but only NHS-A488 labeled centrioles and nucleoli, two biological structures devoid of membranes (Fig. 5B, arrows and arrowheads). The comparison of IgG-A647N, MemExM and NHS-A488 leads us to conclude that IgG-A647N is well retained upon polymerization, homogenization and expansion and represents a convenient tool for membrane labeling in ExM.

**Figure 5:**
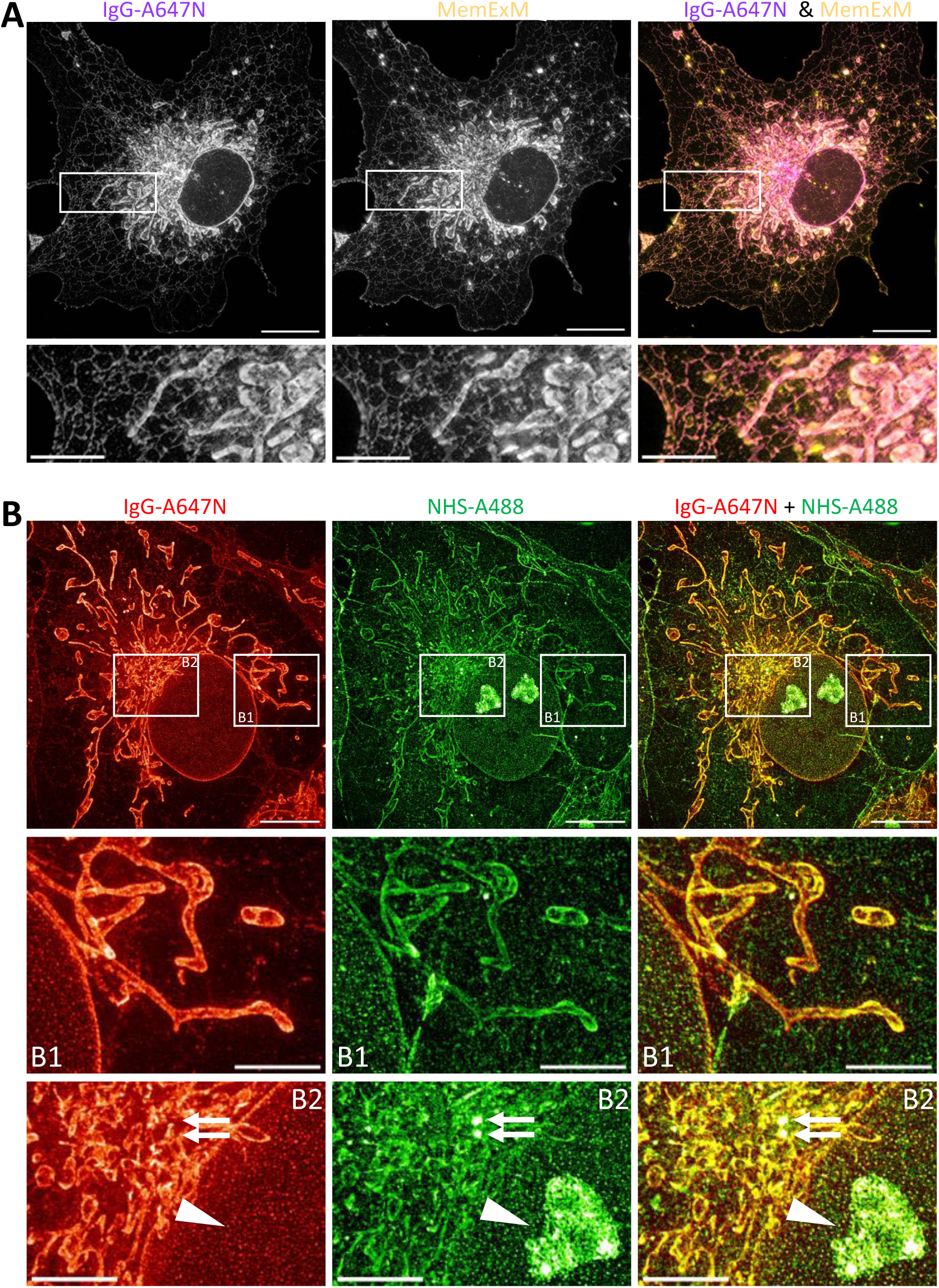
Visualization of cells subjected to Ultrastructure expansion microscopy (U-ExM) and labeled with IgG-A647N, Membrane ExM 561 and/or NHS-coupled ATTO 488. **A:** Fixed and permeabilized COS-7 cells were incubated with IgG-A647N and Membrane ExM 561 (MemExM), subjected to gel-polymerization, expanded in H_2_0 and imaged by wide-field microscopy. Main image and inset reveal highly similar labeling patterns **B**: Fixed and permeabilized COS-7 cells were incubated with IgG-A647N, subjected to gel polymerization, labeled with NHS coupled to ATTO 488 (NHS-A488), expanded in H_2_0 and imaged by wide-field microscopy. IgG-A647N and NHS-A488 displayed with LUT JDM Pop Red and Green, respectively. Main image and insets reveal similar labeling patterns but for centrioles (double arrow) and nucleolus (arrowhead), only labeled by NHS-A488. Bars 10 µm (main figures) and 4 µm (insets).

In a second series of experiments, HeLa cells were fixed, permeabilized, labeled with IgG-A647N, immobilized in a U-ExM gel and denatured with SDS. Gels were then decorated with antibodies against VDAC (an abundant protein of the mitochondrial outer membrane), expanded in water and visualized with a 100X objective (total enlargement ∼560X). The imaging of gels by widefield fluorescence microscopy and 3D-deconvolution revealed VDAC accumulating at the mitochondrial periphery and IgG-A647N molecules labeling inner cristae membranes (Fig. 6A). The density of mitochondrial labeling was remarkable, allowing to image individual cristae and to visualize their regular spacing (Fig. 6A), similar to that observed in living HeLa cells by STED (Liu et al., 2022). In contrast, the non-mitochondrial ER signal was weak and sparse (Fig. 6A). To validate the capacity of IgG-A647N to visualize mitochondrial ultrastructure in ExM, we then analyzed wild-type ρ^+^ cells and mutant ρ° cells devoid of mitochondrial DNA and of functional oxidative phosphorylation complexes. For convenience, we used 143B ρ^+^ and ρ° cells expressing an EGFP protein anchored to the mitochondrial outer membrane (GFPOM, Fig. 3A (Malka et al., 2005)). Cells were fixed, permeabilized, labeled with IgG-A647N and immobilized in a U-ExM gel. To preserve EGFP fluorescence, the U-ExM gel was not homogenized by denaturation (Gambarotto et al., 2021), but by limited proteolysis with proteinase K (Tillberg et al., 2016) and the gel was not expanded in water, but in 5% PBS. The presence of salts in 5% PBS reduces the expansion factor (from 5,6X to 4,1X) but improves the preservation of EGFP-fluorescence (Chen et al., 2021). Imaging with 100X objective (total enlargement ∼410X) revealed that wild-type ρ^+^ cells display filamentous mitochondria that are enveloped by GFPOM and contain cristae membranes labeled with IgG-A647N (Supplementary Fig. S4A). In accordance with electron microscopy analysis (Porteous et al., 1998), mutant ρ° cells contained roundish and swollen mitochondria (labeled with IgG-A647N and GFPOM) that were devoid of inner cristae membranes (Supplementary Fig. S4B).

**Figure 6:**
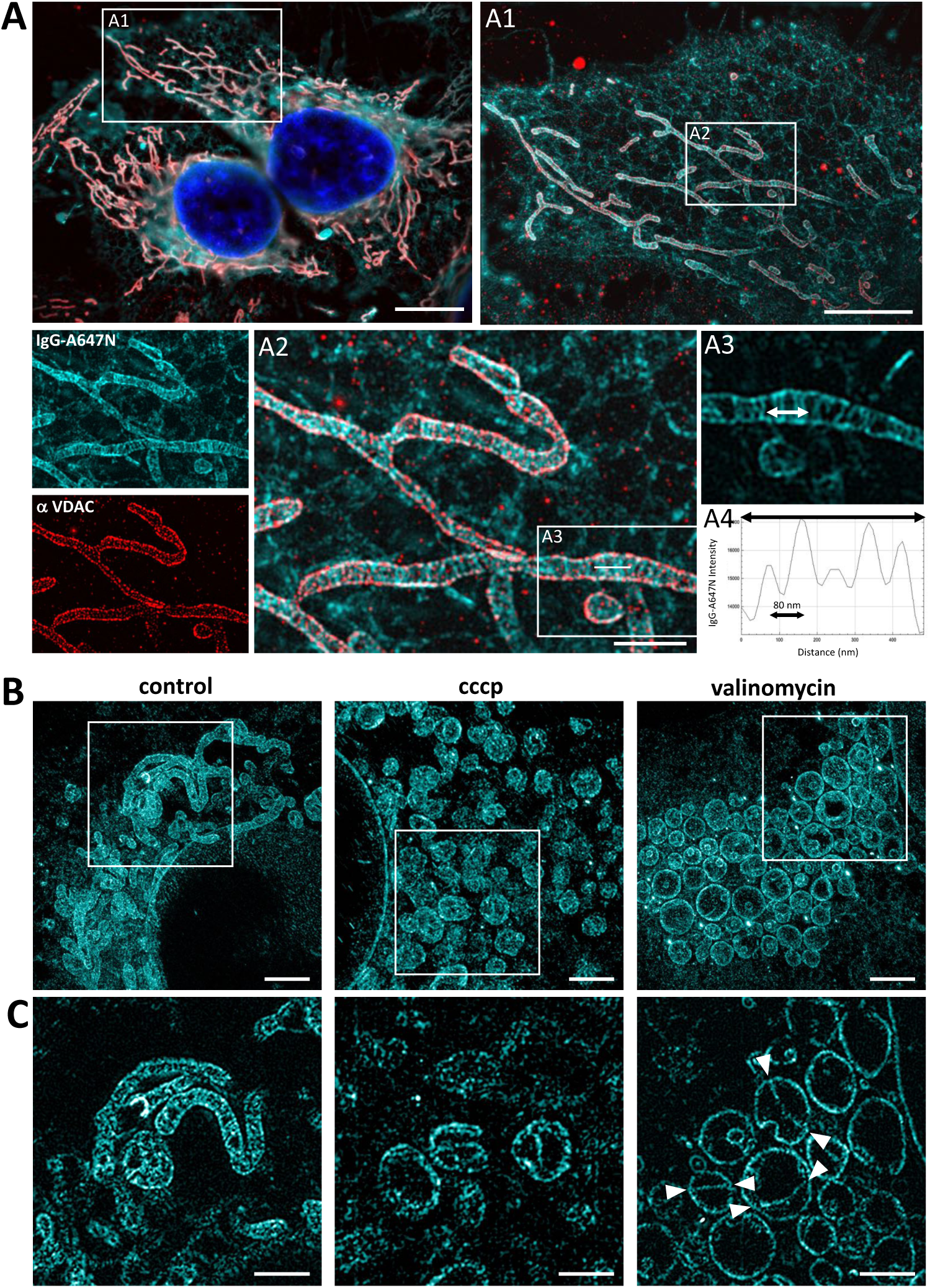
Expansion microscopy (ExM) of cells decorated with IgG-A647N depicts densely labeled mitochondrial membranes and visualizes alterations of mitochondrial ultrastructure. **A:** HeLa cells were fixed, permeabilized, decorated with goat-anti-rabbit IgG-ATTO 647N (IgG-A647N, cyan) and subjected to gel polymerization. Gels were decorated with mouse anti-VDAC and goat-anti-mouse IgG coupled to CF550 (VDAC, red), expanded in H_2_O and visualized by wide-field microscopy. A1 to A3 depict increasingly enlarged insets. A4 displays the intensity profile of cristae membranes along the line drawn in A3. **B, C**: COS-7 cells grown under control conditions or treated with ionophores (cccp or valinomycin) were fixed, permeabilized and decorated with goat anti-rabbit IgG-A647N. Labeled cells were subjected to a first gelation and homogenization before incubation with rabbit anti-goat-IgG-A647N for signal amplification. Gels were then subjected to a neutral and a second expansive gelation, expanded in H_2_O and visualized by wide-field microscopy with 40X (**B**) or 100X objectives (**C**). Images in **C** depict the areas indicated in **B**. Bars: 10 µm (**A**), 5 µm (**A1**), 1 µm (**A2**), 500 nm (**A3**), 5 µm (**B**) and 2 µm (**C**). IgG-A647N displayed with Look-Up Table (LUT) JDM Pop cyan.

Next, we investigated the capacity of IgG-A647N to visualize mitochondrial ultrastructure upon iterative ExM, a protocol achieving 10-20X sample expansion (resulting in 1000X to 8000X dilution of fluorescent label). Having shown that a ‘primary’ label (IgG-A647N from goat) can be decorated with secondary antibodies (donkey anti-goat IgG, supplementary Figure S3), we investigated whether this approach could improve membrane visualization upon at high expansion factors. Fixed and permeabilized cells were labeled with ‘primary’ IgG-A647N, immobilized in a first U-ExM gel, homogenized by denaturation and decorated with ‘secondary’ rabbit anti-goat IgG-A647N. After completion of the iU-ExM protocol (Louvel et al., 2023), reaching 20-fold expansion, samples analyzed with 40X or 100X objectives (Fig. 6B or C, total enlargement ∼800X or 2000X) visualized mitochondrial membranes and ultrastructure. Analysis of COS-7 cells maintained under control conditions, or treated with ionophores dissipating the inner membrane potential (cccp and valinomycin, see Supplementary Fig. S2B), confirmed that mitochondrial depolarization and fragmentation is accompanied by significant ultrastructural changes (Fig. 6B, C), notably the appearance of mitochondrial inner membrane septae (Fig. 6C, arrowheads) across the swollen mitochondria of valinomycin-treated cells that had been observed by electron microscopy in human and in yeast cells (Malka et al., 2005; Sauvanet et al., 2012).

Finally, we applied IgG-A647N to the visualization of the nuclear envelope. It is noteworthy that, although generally represented as a smooth-surfaced outer boundary surrounding the nucleoplasma, the NE is frequently interrupted by invaginations that reach deep within the nucleoplasma as well as by tunnels that traverse it entirely (Malhas et al., 2011). In a majority of reports, imaging of the NE by light microscopy relies on the visualization of the nuclear lamina, a proteinaceous layer between the inner nuclear membrane and the nucleoplasma (Schoen et al., 2017), but recent work has shown that several membrane dyes adapted to ExM directly label the nuclear envelope (Wen et al., 2020; White et al., 2022; Zhuang et al., 2026). Analysis of labeled cells by ExM revealed the capacity of IgG-A647N to achieve dense labeling of the nuclear envelope and of nuclear tunnels (Fig. 7). Nuclear tunnels were observed frequently in U2OS cells (Fig. 7A), but more rarely in COS-7 cells (Fig. 7B). It is tempting to speculate that the latter represent fenestrations appearing at early stages of NE breakdown (Ungricht and Kutay, 2017). Altogether, our experiments demonstrate that high density labeling of mitochondrial and nuclear membranes by IgG-A647N enables their ultrastructural characterization by ExM.

**Figure 7:**
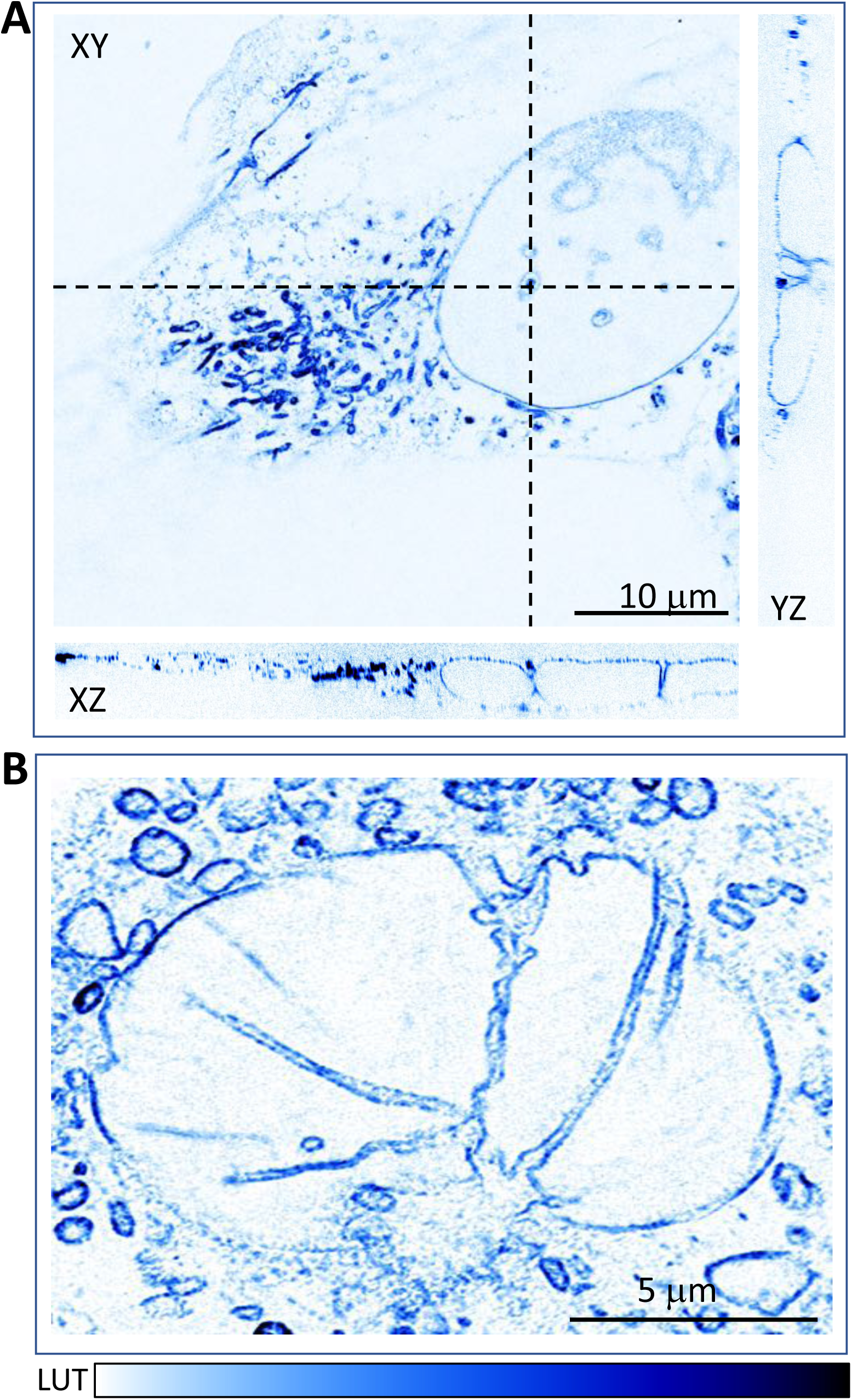
Ultrastructure expansion microscopy (U-ExM) of cells decorated with IgG-A647N depicts densely labeled nuclear envelope membranes, tunnels and invaginations. U2OS cells (**A**) and COS-7 cells (**B**) were fixed, permeabilized, labeled with IgG-A647N, subjected to U-ExM and imaged with ZEISS Apotome 3 (**A**) or wide-field microscopy (**B**). **A:** The dashed lines in XY show the orthogonal YZ and XZ views depicting nuclear tunnels. LUT: JDM Pop Blue inverted.

## Discussion

In this work, we reveal the capacity of IgG molecules coupled to lipophilic ATTO 647N and ATTO 550, as well as of unconjugated hydrophobic dyes (ATTO 647N and Nile Red), to label membranes and organelles of fixed and permeabilized cells. The similar labeling patterns obtained with free and IgG-conjugated dyes, as well as with an established hydrophobic dye (Nile Red), imply that membrane-targeting of IgG molecules is mediated by conjugated lipophilic fluorophores. This conclusion is further supported by the fact that membrane labelling was observed with secondary antibodies with different specificities (anti-mouse, anti-rabbit or anti-rat IgG), generated in different species (goat and rabbit) and provided by different manufacturers (Sigma-Aldrich and Rockland). The fact that membrane labeling was not observed with several IgG-molecules coupled to hydrophilic dyes (ATTO 488, Cy5, AlexaFluor 647 and CF640R, Fig. 1), further confirms that membrane-binding is related to the hydrophobicity and membrane binding capacity of ATTO 647N and ATTO 550 (Hughes et al., 2014).

The observation that secondary antibodies coupled to ATTO 647N produce an unspecific background had been reported (Kolmakov et al., 2010; Wurm et al., 2010), but this is the first study that makes use of this property for labeling of membranes and organelles. The contribution of dye hydrophobicity to membrane labeling agrees with the observation that hydrophobic dyes (i) introduce artefacts in single molecule tracking (Zanetti-Domingues et al., 2013), (ii) influence the labeling properties of reactive fluorescent dye esters (Sheard et al., 2023), (iii) label mitochondria of cultured cells (Han et al., 2017) and (iv) alter the properties of numerous bioconjugates applied *in vivo* (Usama et al., 2021).

We demonstrate that IgG-coupled molecules retain their specific localization in microscopy mounting medium and, most notably, are efficiently anchored within ExM-gels undergoing SDS-mediated denaturation or protease digestion (Figs. 5, 6 and Supplementary Figure S4). Membrane labeling with IgG-A647N or IgG-A550 was relatively fast (15-30 min) and required the permeabilization of cellular membranes with detergents, an obligate step in immunofluorescence experiments (Malka et al., 2007) that allows access of IgGs to intracellular structures while preserving the overall ultrastructure of fixed samples (Humbel et al., 1998). It is tempting to hypothesize that dye-conjugated IgG molecules bind to (the hydrophobic domains of) transmembrane proteins, but it cannot be excluded that dye-coupled IgG (also) bind to detergent-resistant lipids (Lichtenberg et al., 2005). The binding of IgG-coupled dyes to membrane proteins is in accordance with the fact that the most intensely labeled organelles (nuclear envelope and mitochondria) have high protein/lipid ratios (Hahn and Carvalho, 2022; Srere, 1982) and are strongly labeled by fluorescent NHS-esters reacting with proteins (Figure 5B). Of note, a bright labeling of nuclear envelope and mitochondrial network, was also observed with unconjugated hydrophobic dyes (Fig. 2D) and with a fluorescently labelled phospholipid (MemExM, Fig. 5A (Wen et al., 2020)). The labeling of cellular membranes and the preferential labelling of these two organelles has also been observed with a different class of membrane dyes: polycationic peptides that carry a myristoyl chain for membrane anchoring (mCLING, pGk13a) and are applied to fixed and unpermeabilized cells (Damstra et al., 2022) (Shin et al., 2025) (Zhuang et al., 2026). These results indicate that some properties of these stable, convenient and affordable secondary antibodies resemble to those of dedicated membrane-labeling reagents.

The organelles labeled with dye-coupled IgG were identified by different strategies: the nuclear envelope was unambiguously identified by its position and shape (Fig. 2A) and by its immediate vicinity with the DAPI-stained nucleoplasm (Figs. 3B, 4A), mitochondria with MitoTracker^TM^ (Supplementary Fig. S2A) and with mitochondrially targeted fluorescent proteins (Fig 3A) and the ER with specific antibodies and lectins (Fig. 3B). Further studies will be required to establish whether dye-coupled IgG also label other organelles like the Golgi apparatus, endosomes/lysosomes or peroxisomes. It is important to note that the differential labeling intensity of the distinct organelles, notably the stronger labeling of mitochondria, enabled precise identification and selective segmentation of mitochondria with Weka, a trainable segmentation algorithm available in the Fiji platform (Arganda-Carreras et al., 2017). We want to emphasize that the ability to selectively label mitochondria in fixed and permeabilized cells (*i.e.* regardless of their energetic status, supplementary Fig. S2B) is highly advantageous for mitochondrial research, as a majority of mitochondrial dyes are applied *in vivo* and accumulate poorly (or not at all) in depolarized or dysfunctional mitochondria (Poot et al., 1996).

The labeling of the ER by IgG-A647N was visible with conventional (Fig. 3B) and SIM microscopy (Fig. 4A), hardly detectable with STED and ExM (Fig. 4B, 5A, 6A) and invisible with STORM (Supplementary Fig. S3), suggesting that the membrane targets of IgG-A647N are less abundant in ER membranes. In accordance with recent studies (Schroeter et al., 2026), we observed that fluorescently-labeled Concanavalin A (ConA), a lectin binding the mannose residues of core-glycosylated proteins that have not undergone further modification in the Golgi apparatus, is convenient for ER labeling (Fig. 3B). Although the glycans targeted by ConA are known to be enriched in (but not restricted to) the ER (Tartakoff and Vassalli, 1983)(Schroeter et al., 2026), ConA allowed clear visualization of the ER and of mitochondria-ER contacts by SIM and by STED (Fig. 4).

In SIM-images, IgG-A647N revealed a densely labelled membrane pattern resembling that of inner mitochondrial cristae (Fig. 4A) and densely labeled cristae-membranes were clearly visualized by ExM (Fig. 6, Supplementary Fig. S4). With STED, the intramitochondrial distribution of IgG-A647N differed from that of outer membrane Tom20 (Fig. 4B), but the density of mitochondrial labeling was below that observed in STED-studies conducted with small organic vital dyes accumulating in cristae (Liu et al., 2022; Chen et al., 2024). We infer that optimization of STED-parameters may be required to improve visualization of cristae with IgG-A647N. Finally, the ability to label ‘primary’ IgG-A550 with ‘secondary’ antibodies coupled to Alexa Fluor 647 allowed to label mitochondria depicting a cristae-like pattern by STORM microscopy (Supplementary Fig. S3). Of note, STORM microscopy has revealed partially blurred cristae in MitoTracker^TM^-labeled mitochondria (Shim et al., 2012), but visualization of individual cristae was hardly achieved upon detection of inner membrane proteins by immunofluorescence (Klotzsch et al., 2015) (Dlasková et al., 2018) (Palmer et al., 2021). Given the high-density labeling of the inner membrane with ‘primary’ IgG-A647N, we infer that high-resolution imaging of individual cristae membranes by STORM may require further optimization, notably of the density and the blinking of STORM-compatible ‘secondary’ antibodies. Altogether, our results show that IgG-A647N and IgG-A550 are potent tools for visualization of mitochondria and mitochondrial cristae membranes with different techniques of optical super-resolution.

Expansion microscopy being based on the homogenization and expansion of samples subjected to protein-fixatives, the retention of membrane components and the labeling of biological membranes remain challenging. Among the strategies developed for improved membrane-labeling are the improved fixation of lipids (Klimas et al., 2023) and the development of lipid-anchored dyes carrying acryl-groups for their direct anchoring into ExM-gels (Wen et al., 2020) or amino-groups enabling aldehyde-mediated fixation and consequent cross-linking to ExM-gels (Damstra et al., 2022) (Shin et al., 2025; Zhuang et al., 2026). Our results show that IgG-A647N and IgG-A550 bind to membranes of fixed and permeabilized cells and, being protein-based, are cross-linked to ExM-gels by their amino-groups (like mCLING or pGk13a (Damstra et al., 2022) (Shin et al., 2025) (Zhuang et al., 2026)). The almost identical membrane pattern observed with IgG-A647N and with MemExM, (Fig. 5A), indicate that fixation and cross-linking of IgG-A647N is of similar efficacy to the covalent gel-anchoring of MemExM. The dense labeling of mitochondrial cristae with IgG-A647N enabled visualization of mitochondrial ultrastructure (Fig. 6A) and of ultrastructural alterations (Fig. 6B, C; Supplementary Fig. S4) that had been previously characterized by electron microscopy (Malka et al., 2005; Porteous et al., 1998). In addition, the labeling of the nuclear envelope by IgG-A647N and the visualization of its invaginations and tunnels appears highly efficient, and comparable to that achieved with recently developed, ExM-compatible strategies (Wen et al., 2020) (White et al., 2022) (Zhuang et al., 2026). Finally, the ability to amplify the signal of ‘primary’ IgG-A647N with a ‘secondary’ IgG (Fig. 6B, C) appears advantageous for iterative ExM, where high expansion factors lead to significant dilution of fluorescent dyes. In this respect, it is important to note that, by using ‘secondary’ antibodies after the expansion step, the linkage error (10-15 nm (Früh et al., 2021)) is reduced by a factor equal to the expansion factor.

Altogether, our study demonstrates that commercially available secondary antibodies conjugated to lipophilic dyes represent potent and affordable tools for high-density labeling of membranes in conventional, super-resolution and expansion microscopy and that they can be applied for characterization of organellar morphology and ultrastructure. It is tempting to speculate that the coupling of other and/or more lipophilic dyes to IgG molecules (or to other proteins) may represent a general strategy for the generation of novel reagents with improved membrane labeling capacity.

## Materials and Methods

### Reagents

Aequous solutions of paraformaldehyde (32%) and glutaraldehyde (25%) were provided by EMS (Cat # 15714 and 16200). Formaldehyde (FA, 36.5–38%, F8775), Acrylamide (AA, 40%, A4058), N,N′-methylenbisacrylamide (BIS, 2%, M1533), N,N′-(1,2-dihydroxyethylene)-bisacrylamide (DHEBA, 294381), Sodium Acrylate (SA, 97–99%, 408220), Sodium Hydroxyde (NaOH, 206060010), Sodium Dodecyl Sulfate (SDS), Tetramethylethylenediamine (TEMED, 99%, T9281), Poly-L-Lysine (P8920) and Tween20 (P2287) were purchased from Sigma-Aldrich. Ammonium persulfate (APS, EU009-B) and Triton X-100 (ref. 2000-B) were provided by Euromedex and Phosphate Buffered Saline (X0515-500) was purchased from Dutscher.

### Antibodies, lectins and dyes

Anti-mouse IgG and anti-rabbit IgG produced in goat and coupled to ATTO 550 or ATTO 647N were acquired from Sigma-Aldrich (references 40839-1ML-F, 50185-1ML-F or 43394-1ML-F). Rockland anti-rat IgG coupled to ATTO 647N was acquired from antibodies-online (reference ABIN965006). Rabbit-anti-goat coupled to ATTO 647N was provided by Rockland (reference 605-456-013). Goat Anti-Mouse IgG coupled to Cy5 (1,5 mg IgG / ml) was provided by Jackson Immunoresearch (Code Number: 115-175-166). Donkey anti-goat IgG coupled to AlexaFluo 647 was provided by Thermofisher (2 mg IgG / ml, catalogue # # A-21447). Donkey Anti-Goat IgG coupled to CF640R (2 mg IgG / ml) was provided by Biotium (catalogue # 20179). A sample of goat anti-rabbit IgG coupled to Abberior STAR ORANGE was kindly provided by Abberior (STORANGE-1002-500UG). Concanavalin A coupled to AlexaFluor 488 (C21401) and MitoTracker^TM^ Red CMXRos (M7512) were provided by Thermofisher. Wheat Germ Agglutinin coupled to CF488 was provided by Biotium (#29076). Atto 488 NHS ester (reference 41698) and uncoupled ATTO 647N (reference 04507) were acquired from Sigma-Aldrich. Membrane ExM 561 was provided by Chrometra (Membrane ExM -S, 561, Phospholipid). Mouse anti-VDAC1 antibodies were provided by Abcam (ab-4734). Rabbit anti-Tom20 antibodies were provided by Santa Cruz Biotechnology (sc-11415). Anti-calnexin antibodies were provided by Abcam (reference ab22595).

### Cell culture, fixation and permeabilization

The different cell lines (Mouse Embryo Fibroblasts / MEFs, COS-7, HeLa, 143B and U2OS) were grown in complete DMEM with high glucose containing 7% Fetal Bovine Serum. For mitochondrial labeling with MitoTracker^TM^ Red, 100 nM dye was added to the culture medium for 30 min. The 143B and 143B-ρ^0^ cells (Legros et al., 2002) were stably transfected with outer-membrane anchored EGFP as described (Malka et al., 2005). For growth of 143B-ρ^0^ cells, the medium was supplemented with 50 µg/ml uridine. For microscopy, cells grown on glass cover-slips were fixed two to three days after seeding with a mixture of paraformaldehyde/glutaraldehyde (3,2%/0,1%) for 20 min at room temperature. Where indicated, cells were permeabilized with PBS containing 0,1% Triton X-100 for 10 minutes. Further details on culture and immunofluorescence microscopy were as described (Malka et al., 2007) (Barsa et al., 2025).

### Wide-field microscopy, image treatment and image analysis

Cells were imaged with a fully automated Olympus IX81 microscope (Olympus, Tokyo, Japan) equipped with a pE-4000 light source (CoolLED) 40X/1.3NA UPlanFL N, 60X/1.45NA PlanApo and 100×/1.4NA UplanSApo oil immersion objectives (Olympus, Tokyo, Japan), a ORCA Fusion camera (Hamamatsu, Japan) under the control of the CellSENS software (Olympus, Tokyo, Japan). Z-stacks were deconvolved using the ImageJ (Rasband, W.S., ImageJ, U. S. National Institutes of Health, Bethesda, Maryland, USA, https://imagej.net/ij/, 1997-2018), DeconvolutionLab plugin (BIG, EPFL).. Alternatively (Figure 2D, 3B and 7A), cells were imaged with a fully automated Zeiss Axio Imager M2 microscope equipped with a Viluma 7 light source, Plan-Neofluar40X/1.3NA, Plan-Apochromat100X/1.4NA oil immersion objectives, a Axiocam 712 mono R2 camera under the control of the ZEN 3.2 sofware (Carl Zeiss, Iena, Germany). Z stacks (were deconvolved using ZEN 3.2 software using the fast iterative Gold-Meinel algorithm. Images presented are colored selected planes of representative cells.

For structured illumination (Figure 7A), the ApoTome 3 slider was introduced into the field-stop plane of the microscope. 5 positions of the grid were imaged for each z plane. The acquired image stacks were reconstructed and processed according to the manufacturer instructions.

### STED microscopy

Fixed and stained cell samples were imaged with a stimulated emission depletion (STED) super-resolution microscope (Facility Line, Abberior Instruments) using a 60x objective (NA 1.42, oil immersion, UPLXAPO60XO, Olympus). IgG-ATTO 647N and STARORANGE (TOM20) were imaged with pulsed excitation at 640 nm and 561 nm and pulsed depletion at 775 nm, while Alexa Fluor 488 (ConcanvalineA) was imaged with pulsed excitation at 488 nm and pulsed depletion at 595 nm. Detection was adjusted to the spectral ranges 650-755 nm, 571-630 nm and 498-551 nm with time-gating (0.75 ns delay, 8 ns width) employed for STED mode. The pinhole was small for standard avalanche photodiode detection (1 AU for images 1-4, 0.8 AU for images 5-15) and large for MATRIX array-type detection (10.1 AU). The pixel size was fixed at 20 nm, line accumulations ranged from 1-25 repetitions and pixel dwell time was set to 1.5 µs for Alexa 488 and 5 µs for the other channels.

Images were processed via the Lightbox software (Abberior Instruments), employing the TRUESHARP module for MLE deconvolution with background noise suppression. Deconvolution of MATRIX images included a differential detection step for suppression of out-of-focus light. The STAR ORANGE STED channel was pre-processed to remove Alexa Fluor 488 fluorescence excited by the 775 nm STED laser. To this end, the Alexa Fluor 488 channel was smoothed with a Gaussian function, normalized and subtracted from the STAR ORANGE STED channel.

### MI-SIM microscopy

Images were acquired using the Cell Xpanse imaging platform (CSR Biotech, Guangzhou, China), which integrates MI-SIM (Machine Intelligent Structured Illumination Microscopy) with a Yokogawa CSU-W1 spinning disk confocal unit mounted on a Nikon Ti2-E inverted microscope. High-resolution SIM images were obtained using the MI-SIM modality with a 100×/1.49 NA oil immersion objective (Nikon). Samples were illuminated using 405 nm, 488 nm, 561 nm, and 638 nm lasers (300 mW). Z-stack images were acquired from bottom to top with step size of 0.100 µm, total 19 slices. CSR Biotech MicroscopeX FINER software used for image post processing. SIM images were collected and analyzed as described previously (Huang et al., 2018).

### STORM microscopy

For STORM microscopy, cells labeled with Goat anti-mouse IgG coupled to Atto550 were labeled with a Donkey anti Goat IgG coupled to AlexaFluor 647. 2D single color dSTORM images were acquired with a SAFe MN360 module and Nexus laser combiner (Abbelight, France). The Abbelight system was mounted on an Evident IX83 microscope equipped with a 100×/1.5 NA TIRF objective and a ZDC830 maintenance focus system. The samples were illuminated with 640nm laser (1W, Cobolt) set at 80% over a large field of view up to 150×150um using Abbelight’s ASTER illumination. Short pulses of 405 nm laser (1 to 5% ramp, 50 mW, Oxxius laser) was used for the reactivation of the fluorophores while maintaining single molecule regime. Laser illumination was set to obtain a highly inclined laminated optical sheet (HiLo) illumination mode using Abbelight automated TIRF calibration. Images were acquired using a sCMOS Hamamatsu Fusion camera (Hamamatsu, Japan) at an exposure time of 20ms. The NEO_LiveImaging software (Abbelight) was used to drive the acquisition and NEO_Analysis (Abbelight) was used to localize particles, correct the drift and, finally, to reconstruct the final images with localization filtered by localization precision (max. 30nm).

### Expansion Microscopy

Fixed cells were incubated for 3 h in anchoring solution (1% AA, 0.7% FA in 1× PBS) at 37 °C, washed in PBS and processed for gelation in a modified U-ExM monomer solution (10% AA, 19% SA, 0.025% BIS in 1× PBS) containing 0.5% TEMED and APS. Next, cells were incubated for 5 min on ice followed by 1 h at 37 °C. Gels were hydrated in denaturation buffer (200 mM SDS, 50 mM Tris pH 6.8) and incubated for 1 h at 73 °C in. Gels were washed with PBS and allowed to expand by successive washing in milliQ-H2O until complete expansion.

For immunolabeling, expanded gels were shrunk back by two successive washes in PBS for 15 minutes each and incubated with primary antibodies in 1X PBS/2%BSA ON at 37°C. Gels were then washed 3 times with 1XPBS/0.1%Tween-20 for 20 minutes each and incubated secondary antibodies in 1X PBS/2%BSA 3h at 37°C. Gels were next allowed to re-expand in milliQ-H2O before imaging.

Iterative ExM was processed as described (M’Saad and Bewersdorf, 2020; Louvel et al., 2023). Briefly, fixed cells were incubated with fixation in 0.7% formaldehyde + 1% acrylamide (w/v) in 1× PBS for 3 h at 37 °C. After washing the cells in PBS, coverslips were mounted in gelation chambers constituted by a microscopy slide bordered by 2 stacks of 0.17mm thick 22×22mm coverslips bordering the cells containing coverslip and covered by a 22×22mm coverslip. Cells were incubated in the monomer solution containing (19% (w/v) sodium acrylate (SA) + 10% acrylamide (AA) (w/v) + 0.1% (w/v) DHEBA (N,N′-(1,2-dihydroxyethylene) bisacrylamide) + 0.25% (v/v) TEMED (N,N,N′,N′-tetramethylethylenediamine) + 0.25% (w/v) Ammonium persulfate (APS) in PBS and incubated for 1 h at 37 °C in a humid chamber to allow polymerization. Next, the gelation chambers were carefully dismantled and the coverslips/gels were dipped in 2 mL denaturation buffer in a 6 well plate under shaking until the gel detached from the coverslip. The gels were next transferred in a 1.5 mL Eppendorf tube with 1 mL of fresh denaturation buffer and incubated for 1 h at 73 °C. Then, the gels were expanded in milliQ-H2O with at least 3 washes, until the expansion of the gel plateaus. Next, the gels were cut into a 1 cm2 piece and embedded in a neutral gel (10% AA; 0.05% DHEBA; 0.05% APS/TEMED in ddH2O). Embedded gels were incubated in a third monomer solution containing (19% (w/v) SA + 10% AA (w/v) + 0.1% (w/v) BIS + 0.05% (v/v) TEMED + 0.05% (w/v) APS in PBS). After polymerization, the first gel containing DHEBA crosslinkers was dissolved by incubating it in 0.2 M NaOH for 1 h. After several washes, the gels were allowed to expand in milliQ water until final expansion.

Pan-Staining was processed as described (M’Saad and Bewersdorf, 2020). Briefly, after final expansion in water, gels were incubated in 100 mM sodium bicarbonate containing 20 ug/ml NHSester Atto488 for 1.5 h at RT. Gels were subsequently washed in PBS/Tween20 0.1%, 3 to 5 times for 20 min each, and allowed to expand in milliQ water until final expansion.

## Acknowledgements

We are indebted to Abberior and Gero Schlötel for assistance in acquisition and treatment of STED images as well as Abbelight and Benjamin Compans for assistance in acquisition and treatment of STORM images. We thank CSR Europe GmbH and Ipek Bonnet for assistance in acquisition and treatment of MI-SIM images by using their commercial super-resolution microscope Cell Xpanse, data acquisition, SR image reconstruction, analysis and discussion. We thank Bordeaux Imaging Center (a service unit of the CNRS, INSERM and Bordeaux University, member of the national infrastructure France BioImaging supported by the French National Research Agency / ANR-24-INBS-0005 FBI BIOGEN) for organizing workshops and demonstrations allowing us to use STED, STORM and MI-SIM microscopy. We are grateful to Eloïse Bertiaux, Giulia Bertolin, Ingrid Chamma, Mayeul Collot, Eloina Corradi, Magali Grison and Sandrine Pouvreau for careful reading of the manuscript and for critical comments and suggestions. We thank Mónica Fernández-Monreal for sharing reagents and thoughts. We used the HeLa cell line that was established from the tumor cells of Henrietta Lacks in 1951. We are grateful to Henrietta Lacks, now deceased, and to her surviving family members for their contributions to biomedical research and to the progress of this project. This work was supported by grants from AFM (28981 to M.R.) and ANR (ANR-22-CE14-0040 to A.M.).

## Author contributions

J.D. and M.R. initiated, conceived and supervised the project. J.D., S.P.P, L.H. and M.R. performed and analyzed experiments. A.M. and A.D. supported the project. M.R. wrote the original draft. All authors reviewed and edited the manuscript.

## References

Aknine, N., R. Pelletier, and A.S. Klymchenko. 2025. Lipid-Directed Covalent Labeling of Plasma Membranes for Long-Term Imaging, Barcoding and Manipulation of Cells. JACS Au. 5:922–936. doi:10.1021/jacsau.4c01134.

Arganda-Carreras, I., V. Kaynig, C. Rueden, K.W. Eliceiri, J. Schindelin, A. Cardona, and H. Sebastian Seung. 2017. Trainable Weka Segmentation: a machine learning tool for microscopy pixel classification. Bioinformatics. 33:2424–2426. doi:10.1093/bioinformatics/btx180.

Barsa, C., J. Perrin, C. David, A. Mourier, and M. Rojo. 2025. A cellular assay to determine the fusion capacity of MFN2 variants linked to Charcot-Marie-Tooth disease of type 2 A. Sci Rep. 15:9971. doi:10.1038/s41598-025-93702-1.

Chen, J., T. Stephan, F. Gaedke, T. Liu, Y. Li, A. Schauss, P. Chen, V. Wulff, S. Jakobs, C. Jüngst, and Z. Chen. 2024. An aldehyde-crosslinking mitochondrial probe for STED imaging in fixed cells. Proceedings of the National Academy of Sciences. 121:e2317703121. doi:10.1073/pnas.2317703121.

Chen, R., X. Cheng, Y. Zhang, X. Yang, Y. Wang, X. Liu, and S. Zeng. 2021. Expansion tomography for large volume tissue imaging with nanoscale resolution. Biomed Opt Express. 12:5614–5628. doi:10.1364/BOE.431696.

Collot, M., P. Ashokkumar, H. Anton, E. Boutant, O. Faklaris, T. Galli, Y. Mély, L. Danglot, and A.S. Klymchenko. 2019. MemBright: A Family of Fluorescent Membrane Probes for Advanced Cellular Imaging and Neuroscience. Cell Chemical Biology. 26:600–614.e7. doi:10.1016/j.chembiol.2019.01.009.

Damstra, H.G.J., B. Mohar, M. Eddison, A. Akhmanova, L.C. Kapitein, and P.W. Tillberg. 2022. Visualizing cellular and tissue ultrastructure using Ten-fold Robust Expansion Microscopy (TREx). Elife. 11. doi:10.7554/eLife.73775.

Dempsey, G.T., J.C. Vaughan, K.H. Chen, M. Bates, and X. Zhuang. 2011. Evaluation of fluorophores for optimal performance in localization-based super-resolution imaging. Nat Meth. 8:1027–1036. doi:10.1038/nmeth.1768.

Dlasková, A., H. Engstová, T. Špaček, A. Kahancová, V. Pavluch, K. Smolková, J. Špačková, M. Bartoš, L.P. Hlavatá, and P. Ježek. 2018. 3D super-resolution microscopy reflects mitochondrial cristae alternations and mtDNA nucleoid size and distribution. BBA - Bioenergetics. 1859:829–844. doi:10.1016/j.bbabio.2018.04.013.

Fam, T.K., A.S. Klymchenko, and M. Collot. 2018. Recent Advances in Fluorescent Probes for Lipid Droplets. Materials (Basel). 11. doi:10.3390/ma11091768.

Freundt, E.C., M. Czapiga, and M.J. Lenardo. 2007. Photoconversion of Lysotracker Red to a green fluorescent molecule. Cell Res. 17:956–958. doi:10.1038/cr.2007.80.

Früh, S.M., U. Matti, P.R. Spycher, M. Rubini, S. Lickert, T. Schlichthaerle, R. Jungmann, V. Vogel, J. Ries, and I. Schoen. 2021. Site-Specifically-Labeled Antibodies for Super-Resolution Microscopy Reveal In Situ Linkage Errors. ACS Nano. 15:12161–12170. doi:10.1021/acsnano.1c03677.

Gambarotto, D., V. Hamel, and P. Guichard. 2021. Ultrastructure expansion microscopy (U-ExM). Methods Cell Biol. 161:57–81. doi:10.1016/bs.mcb.2020.05.006.

Guillery, O., F. Malka, P. Frachon, M. Rojo, and A. Lombès. 2008a. Modulation of mitochondrial morphology by bioenergetics defects in primary human fibroblasts. Neuromuscul Disord. 18:319–330. doi:10.1016/j.nmd.2007.12.008.

Guillery, O., F. Malka, T. Landes, E. Guillou, C. Blackstone, A. Lombès, P. Belenguer, D. Arnoult, and M. Rojo. 2008b. Metalloprotease-mediated OPA1 processing is modulated by the mitochondrial membrane potential. Biol Cell. 100:315–325. doi:10.1042/BC20070110.

Hahn, L., and P. Carvalho. 2022. Making and breaking the inner nuclear membrane proteome. Curr Opin Cell Biol. 78:102115. doi:10.1016/j.ceb.2022.102115.

Han, Y., M. Li, F. Qiu, M. Zhang, and Y.-H. Zhang. 2017. Cell-permeable organic fluorescent probes for live-cell long-term super-resolution imaging reveal lysosome-mitochondrion interactions. Nat Comms. 8:1307. doi:10.1038/s41467-017-01503-6.

Hemel, I.M.G.M., B.P.H. Engelen, N. Luber, and M. Gerards. 2021. A hitchhiker’s guide to mitochondrial quantification. MITOCHONDRION. 59:216–224. doi:10.1016/j.mito.2021.06.005.

Hughes, L.D., R.J. Rawle, and S.G. Boxer. 2014. Choose your label wisely: water-soluble fluorophores often interact with lipid bilayers. PLoS ONE. 9:e87649. doi:10.1371/journal.pone.0087649.

Humbel, B.M., M.D. de Jong, W.H. Müller, and A.J. Verkleij. 1998. Pre-embedding immunolabeling for electron microscopy: an evaluation of permeabilization methods and markers. Microsc Res Tech. 42:43–58. doi:10.1002/(SICI)1097-0029(19980701)42:1<43::AID-JEMT6>3.0.CO;2-S.

Hümpfer, N., R. Thielhorn, and H. Ewers. 2024. Expanding boundaries - a cell biologist’s guide to expansion microscopy. Journal of Cell Science. 137. doi:10.1242/jcs.260765.

Kim, S.Y., K. Strucinska, B. Osei, K. Han, S.-K. Kwon, and T.L. Lewis. 2022. Neuronal mitochondrial morphology is significantly affected by both fixative and oxygen level during perfusion. Front Mol Neurosci. 15:1042616. doi:10.3389/fnmol.2022.1042616.

Klimas, A., B.R. Gallagher, P. Wijesekara, S. Fekir, E.F. DiBernardo, Z. Cheng, D.B. Stolz, F. Cambi, S.C. Watkins, S.L. Brody, A. Horani, A.L. Barth, C.I. Moore, X. Ren, and Y. Zhao. 2023. Magnify is a universal molecular anchoring strategy for expansion microscopy. Nat Biotech. doi:10.1038/s41587-022-01546-1.

Klotzsch, E., A. Smorodchenko, L. Löfler, R. Moldzio, E. Parkinson, G.J. Schütz, and E.E. Pohl. 2015. Superresolution microscopy reveals spatial separation of UCP4 and F0F1-ATP synthase in neuronal mitochondria. Proceedings of the National Academy of Sciences. 112:130–135. doi:10.1073/pnas.1415261112.

Kolmakov, K., V.N. Belov, J. Bierwagen, C. Ringemann, V. Müller, C. Eggeling, and S.W. Hell. 2010. Red-emitting rhodamine dyes for fluorescence microscopy and nanoscopy. Chemistry. 16:158–166. doi:10.1002/chem.200902309.

Lambert, T.J., and J.C. Waters. 2017. Navigating challenges in the application of superresolution microscopy. The Journal of Cell Biology. 216:53–63. doi:10.1083/jcb.201610011.

Laporte, M.H., N. Klena, V. Hamel, and P. Guichard. 2022. Visualizing the native cellular organization by coupling cryofixation with expansion microscopy (Cryo-ExM). Nat Meth. 19:216–222. doi:10.1038/s41592-021-01356-4.

Lefebvre, A.E.Y.T., G. Sturm, T.-Y. Lin, E. Stoops, M.P. López, B. Kaufmann-Malaga, and K. Hake. 2025. Nellie: automated organelle segmentation, tracking and hierarchical feature extraction in 2D/3D live-cell microscopy. Nat Meth. 22:751–763. doi:10.1038/s41592-025-02612-7.

Legros, F., A. Lombès, P. Frachon, and M. Rojo. 2002. Mitochondrial fusion in human cells is efficient, requires the inner membrane potential, and is mediated by mitofusins. Mol Biol Cell. 13:4343–4354. doi:10.1091/mbc.E02-06-0330.

Lichtenberg, D., F.M. Goñi, and H. Heerklotz. 2005. Detergent-resistant membranes should not be identified with membrane rafts. Trends in Biochemical Sciences. 30:430–436. doi:10.1016/j.tibs.2005.06.004.

Liu, T., T. Stephan, P. Chen, J. Keller-Findeisen, J. Chen, D. Riedel, Z. Yang, S. Jakobs, and Z. Chen. 2022. Multi-color live-cell STED nanoscopy of mitochondria with a gentle inner membrane stain. Proceedings of the National Academy of Sciences. 119:e2215799119. doi:10.1073/pnas.2215799119.

Louvel, V., R. Haase, O. Mercey, M.H. Laporte, T. Eloy, É. Baudrier, D. Fortun, D. Soldati-Favre, V. Hamel, and P. Guichard. 2023. iU-ExM: nanoscopy of organelles and tissues with iterative ultrastructure expansion microscopy. Nat Comms. 14:7893. doi:10.1038/s41467-023-43582-8.

Malhas, A., C. Goulbourne, and D.J. Vaux. 2011. The nucleoplasmic reticulum: form and function. Trends Cell Biol. 21:362–373. doi:10.1016/j.tcb.2011.03.008.

Malka, F., K. Auré, S. Goffart, J.N. Spelbrink, and M. Rojo. 2007. The mitochondria of cultured mammalian cells: I. Analysis by immunofluorescence microscopy, histochemistry, subcellular fractionation, and cell fusion. Methods Mol Biol. 372:3–16.

Malka, F., O. Guillery, C. Cifuentes-Diaz, E. Guillou, P. Belenguer, A. Lombès, and M. Rojo. 2005. Separate fusion of outer and inner mitochondrial membranes. EMBO Rep. 6:853–859. doi:10.1038/sj.embor.7400488.

Marin, Z., L.A. Fuentes, J. Bewersdorf, and D. Baddeley. 2023. Extracting nanoscale membrane morphology from single-molecule localizations. Biophys J. 122:3022–3030. doi:10.1016/j.bpj.2023.06.010.

Mezache, L., and C. Leterrier. 2025. Advancing Super-Resolution Microscopy: Recent Innovations in Commercial Instruments. Microsc Microanal. 31. doi:10.1093/mam/ozaf004.

M’Saad, O., and J. Bewersdorf. 2020. Light microscopy of proteins in their ultrastructural context. Nat Comms. 11:3850. doi:10.1038/s41467-020-17523-8.

Palmer, C.S., J. Lou, B. Kouskousis, E. Pandzic, A.J. Anderson, Y. Kang, E. Hinde, and D. Stojanovski. 2021. Super-resolution microscopy reveals the arrangement of inner membrane protein complexes in mammalian mitochondria. Journal of Cell Science. 134. doi:10.1242/jcs.252197.

Poot, M., Y.Z. Zhang, J.A. Krämer, K.S. Wells, L.J. Jones, D.K. Hanzel, A.G. Lugade, V.L. Singer, and R.P. Haugland. 1996. Analysis of mitochondrial morphology and function with novel fixable fluorescent stains. Journal of Histochemistry and Cytochemistry. 44:1363–1372.

Porteous, W.K., A.M. James, P.W. Sheard, C.M. Porteous, M.A. Packer, S.J. Hyslop, J.V. Melton, C.Y. Pang, Y.H. Wei, and M.P. Murphy. 1998. Bioenergetic consequences of accumulating the common 4977-bp mitochondrial DNA deletion. Eur J Biochem. 257:192–201. doi:10.1046/j.1432-1327.1998.2570192.x.

Revelo, N.H., D. Kamin, S. Truckenbrodt, A.B. Wong, K. Reuter-Jessen, E. Reisinger, T. Moser, and S.O. Rizzoli. 2014. A new probe for super-resolution imaging of membranes elucidates trafficking pathways. The Journal of Cell Biology. 205:591–606. doi:10.1083/jcb.201402066.

Sauvanet, C., S. Duvezin-Caubet, B. Salin, C. David, A. Massoni-Laporte, J.-P. Di Rago, and M. Rojo. 2012. Mitochondrial DNA mutations provoke dominant inhibition of mitochondrial inner membrane fusion. PLoS ONE. 7:e49639. doi:10.1371/journal.pone.0049639.

Schoen, I., L. Aires, J. Ries, and V. Vogel. 2017. Nanoscale invaginations of the nuclear envelope: Shedding new light on wormholes with elusive function. Nucleus. 8:506–514. doi:10.1080/19491034.2017.1337621.

Schroeter, H.G., S. Sass, M. Heilemann, T. Kuner, and M. Klevanski. 2026. Nanoscale Mapping of the Subcellular Glycosylation Landscape. Adv Sci (Weinh). 13:e06731. doi:10.1002/advs.202506731.

Sheard, T.M.D., T.B. Shakespeare, R.S. Seehra, M.E. Spencer, K.M. Suen, and I. Jayasinghe. 2023. Differential labelling of human sub-cellular compartments with fluorescent dye esters and expansion microscopy. Nanoscale. 15:18489–18499. doi:10.1039/d3nr01129a.

Shim, S.-H., C. Xia, G. Zhong, H.P. Babcock, J.C. Vaughan, B. Huang, X. Wang, C. Xu, G.-Q. Bi, and X. Zhuang. 2012. Super-resolution fluorescence imaging of organelles in live cells with photoswitchable membrane probes. Proceedings of the National Academy of Sciences. 109:13978–13983. doi:10.1073/pnas.1201882109.

Shin, T.W., H. Wang, C. Zhang, B. An, Y. Lu, E. Zhang, X. Lu, E.D. Karagiannis, J.S. Kang, A. Emenari, P. Symvoulidis, S. Asano, L. Lin, E.K. Costa, IMAXT Grand Challenge Consortium, A.H. Marblestone, N. Kasthuri, L.-H. Tsai, and E.S. Boyden. 2025. Dense, continuous membrane labeling and expansion microscopy visualization of ultrastructure in tissues. Nat Comms. 16:1579. doi:10.1038/s41467-025-56641-z.

Snapp, E.L., R.S. Hegde, M. Francolini, F. Lombardo, S. Colombo, E. Pedrazzini, N. Borgese, and J. Lippincott-Schwartz. 2003. Formation of stacked ER cisternae by low affinity protein interactions. J Cell Biol. 163:257–269. doi:10.1083/jcb.200306020.

Srere, P.A. 1982. The structure of the mitochondrial inner membrane-matrix compartment. Trends in Biochemical Sciences. 7:375–378. doi:10.1016/0968-0004(82)90119-0.

Stoldt, S., T. Stephan, D.C. Jans, C. Brüser, F. Lange, J. Keller-Findeisen, D. Riedel, S.W. Hell, and S. Jakobs. 2019. Mic60 exhibits a coordinated clustered distribution along and across yeast and mammalian mitochondria. Proceedings of the National Academy of Sciences. doi:10.1073/pnas.1820364116.

Swaisgood, M., and M. Schindler. 1989. Lateral diffusion of lectin receptors in fibroblast membranes as a function of cell shape. Experimental Cell Research. 180:515–528. doi:10.1016/0014-4827(89)90078-5.

Tartakoff, A.M., and P. Vassalli. 1983. Lectin-binding sites as markers of Golgi subcompartments: proximal-to-distal maturation of oligosaccharides. J Cell Biol. 97:1243–1248. doi:10.1083/jcb.97.4.1243.

Tillberg, P.W., F. Chen, K.D. Piatkevich, Y. Zhao, C.-C.J. Yu, B.P. English, L. Gao, A. Martorell, H.-J. Suk, F. Yoshida, E.M. DeGennaro, D.H. Roossien, G. Gong, U. Seneviratne, S.R. Tannenbaum, R. Desimone, D. Cai, and E.S. Boyden. 2016. Protein-retention expansion microscopy of cells and tissues labeled using standard fluorescent proteins and antibodies. Nat Biotech. 34:987–992. doi:10.1038/nbt.3625.

Tondera, D., S. Grandemange, A. Jourdain, M. Karbowski, Y. Mattenberger, S. Herzig, S. Da Cruz, P. Clerc, I. Raschke, C. Merkwirth, S. Ehses, F. Krause, D.C. Chan, C. Alexander, C. Bauer, R. Youle, T. Langer, and J.-C. Martinou. 2009. SLP-2 is required for stress-induced mitochondrial hyperfusion. EMBO J. 28:1589–1600. doi:10.1038/emboj.2009.89.

Ungricht, R., and U. Kutay. 2017. Mechanisms and functions of nuclear envelope remodelling. Nat Rev Mol Cell Biol. 18:229–245. doi:10.1038/nrm.2016.153.

Usama, S.M., E.R. Thapaliya, M.P. Luciano, and M.J. Schnermann. 2021. Not so innocent: Impact of fluorophore chemistry on the in vivo properties of bioconjugates. Curr Opin Chem Biol. 63:38–45. doi:10.1016/j.cbpa.2021.01.009.

Wen, G., M. Vanheusden, A. Acke, D. Valli, R.K. Neely, V. Leen, and J. Hofkens. 2020. Evaluation of Direct Grafting Strategies via Trivalent Anchoring for Enabling Lipid Membrane and Cytoskeleton Staining in Expansion Microscopy. ACS Nano. doi:10.1021/acsnano.9b09259.

White, B.M., P. Kumar, A.N. Conwell, K. Wu, and J.M. Baskin. 2022. Lipid Expansion Microscopy. J Am Chem Soc. 144:18212–18217. doi:10.1021/jacs.2c03743.

Wurm, C.A., D. Neumann, R. Schmidt, A. Egner, and S. Jakobs. 2010. Sample preparation for STED microscopy. Methods Mol Biol. 591:185–199. doi:10.1007/978-1-60761-404-3_11.

Zanetti-Domingues, L.C., C.J. Tynan, D.J. Rolfe, D.T. Clarke, and M. Martin-Fernandez. 2013. Hydrophobic fluorescent probes introduce artifacts into single molecule tracking experiments due to non-specific binding. PLoS ONE. 8:e74200. doi:10.1371/journal.pone.0074200.

Zhuang, Y., Z. Zhang, Z. Dai, and X. Shi. 2026. Landscape expansion microscopy reveals interactions between membrane and phase-separated organelles. The Journal of Cell Biology. 225. doi:10.1083/jcb.202502035.

