## Supplemental Material for "From background to foreground: secondary antibodies coupled to lipophilic ATTO dyes enable high-density membrane labeling in super-resolution and expansion microscopy"

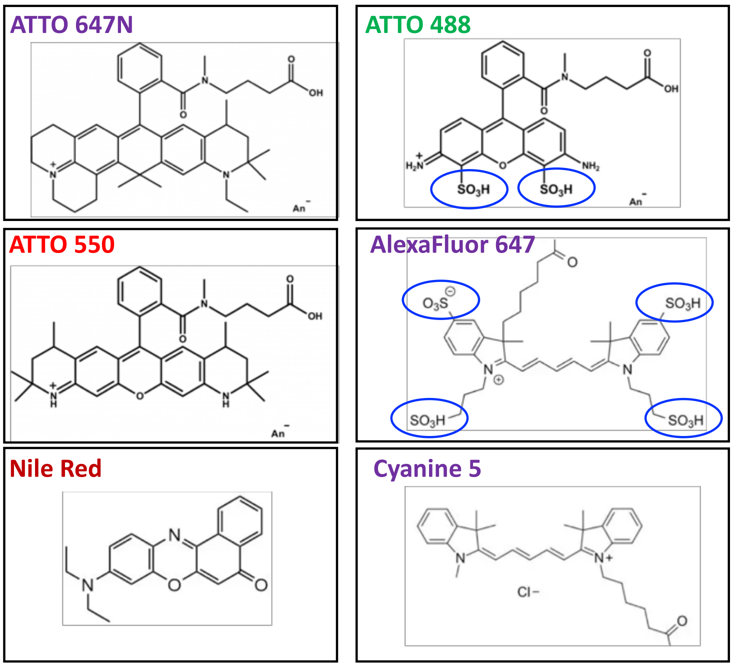


**Supplementary Figure S1: Molecular structure of selected fluorescent dyes.** Left column: lipophilic ATTO 647N, ATTO 550 and Nile Red. Right column: hydrophilic ATTO 488, AlexaFluor 647 and cyanine 5. Polar sulfate groups are highlighted.

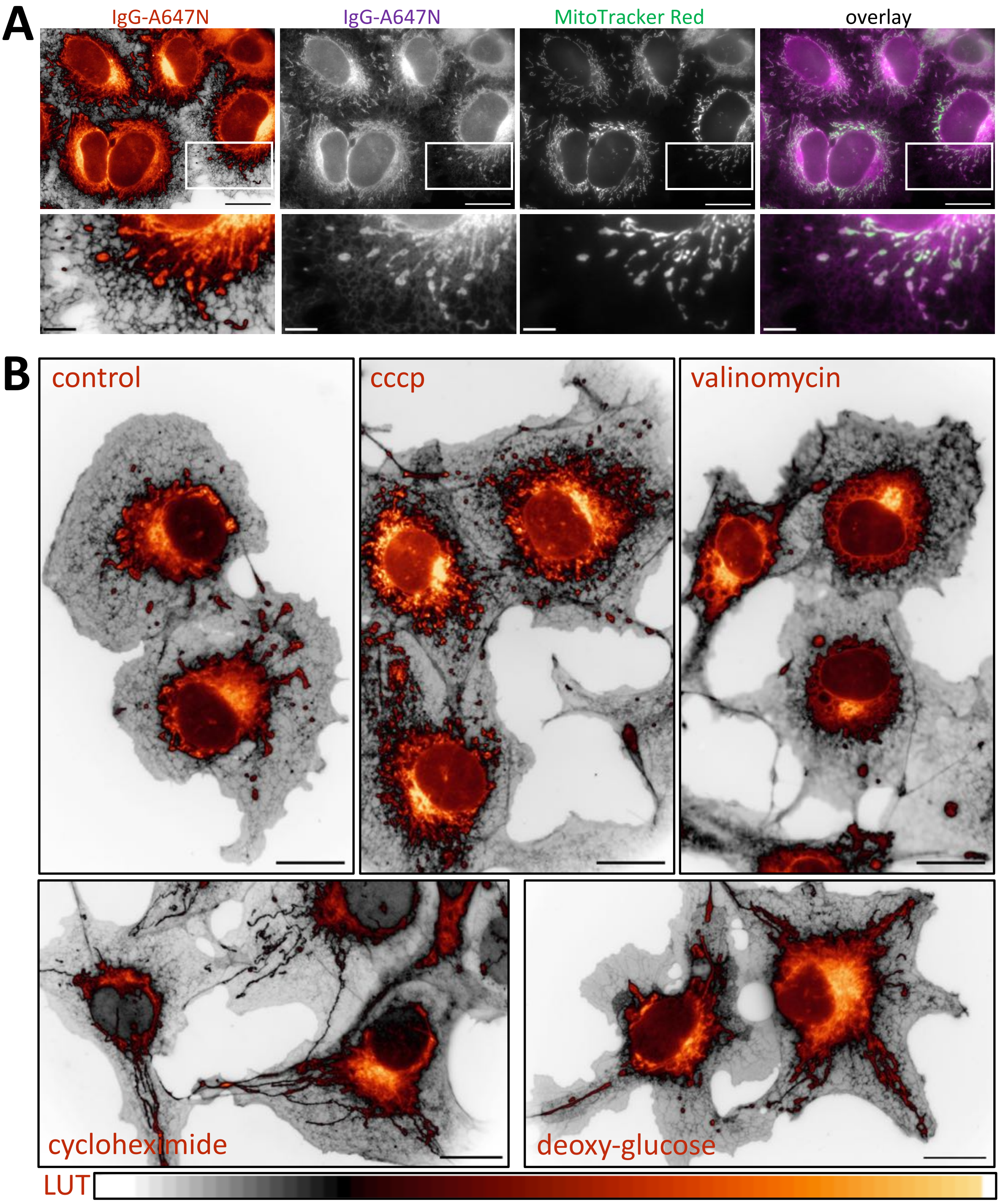


**Supplementary Figure S2: Imaging of mitochondria with IgG-A647N and Mitotracker$^{TM}$ red. A:** Hela cells labeled *in vivo* with Mitotracker$^{TM}$ red were fixed, permeabilized and labeled with IgG-A647N. IgG-labeling is shown with the indicated LUT, in grey levels or in overlay with Mitotracker red. For optimal overlay, IgG-A647N and Mitotracker Red are shown in magenta and green, respectively. **B:** COS-7 grown under control conditions or treated for 4 hours with cccp, valinomycin, cycloheximide or deoxy-glucose were fixed, permeabilized, decorated with IgG-A647N and imaged by wide-field microscopy. Images treated with the indicated LUT depict depolarized fragmented (cccp) or swollen and perinuclearly clustered mitochondria (valinomycin). Cells treated with cycloheximide or deoxy-glucose depict filamentous mitochondria. Bars 20 µm (main images) or 5 µm (insets). LUT: MQ div-autumn.

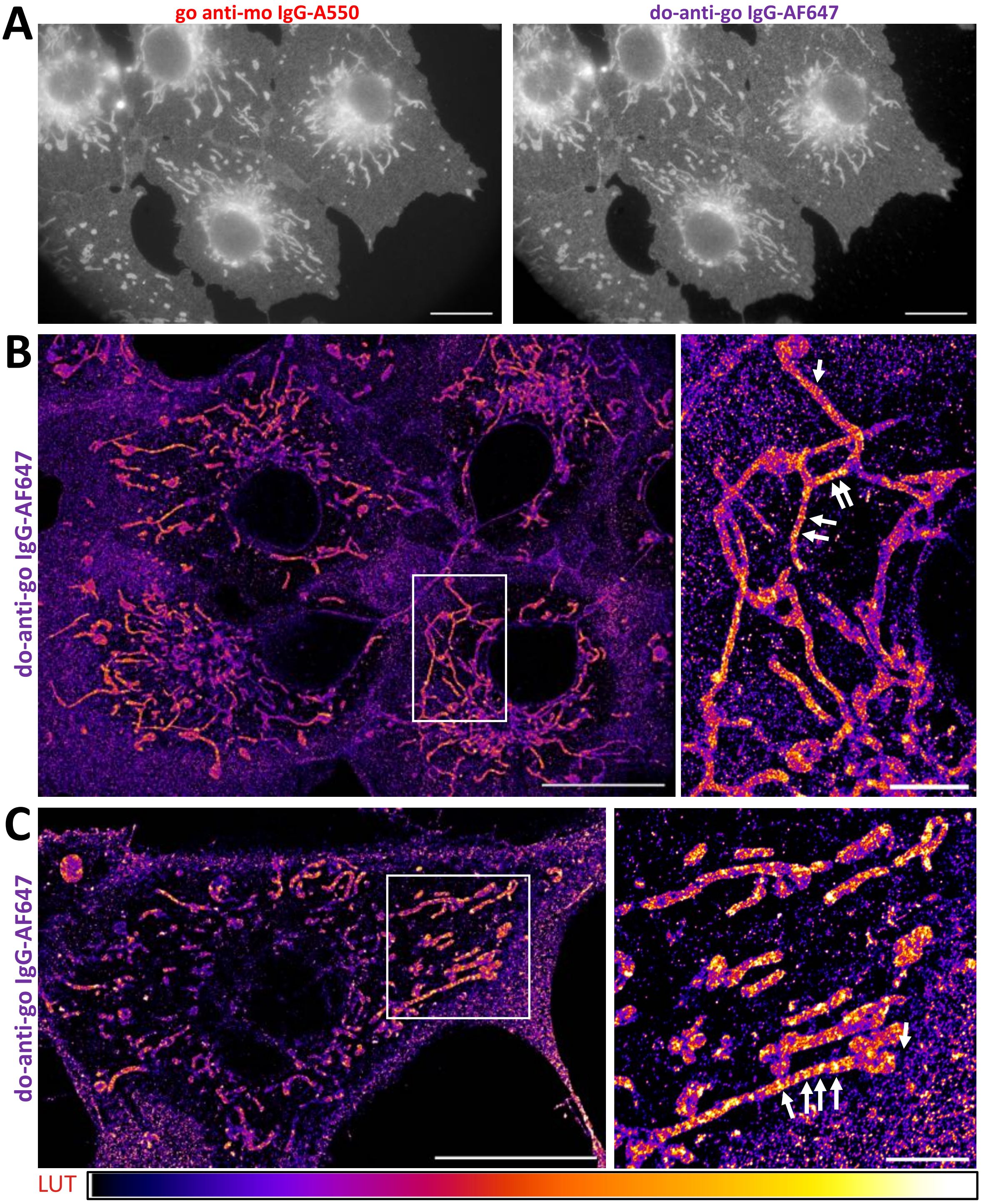


**Supplementary Figure S3: Membranes primarily labeled with goat-IgG-A550 can be visualized by STORM microscopy with secondary donkey anti-goat-IgG coupled to AlexaFluor 647.** COS-7 were fixed, permeabilized and successively decorated with go-IgG-A550 and with do-anti-go-IgG-AlexaFluor 647. **A**: Primary IgG-A550 and secondary IgG-AlexaFluor 647 imaged by wide-field fluorescence microscopy. **B, C**: Secondary IgG AlexaFluor 647 visualized by STORM. Images pseudocolored with the Fire Look-Up Table (LUT). Bars 20 µm (main figures, left) and 4 µm (enlarged insets, right). Arrows point to cristae-like structures.

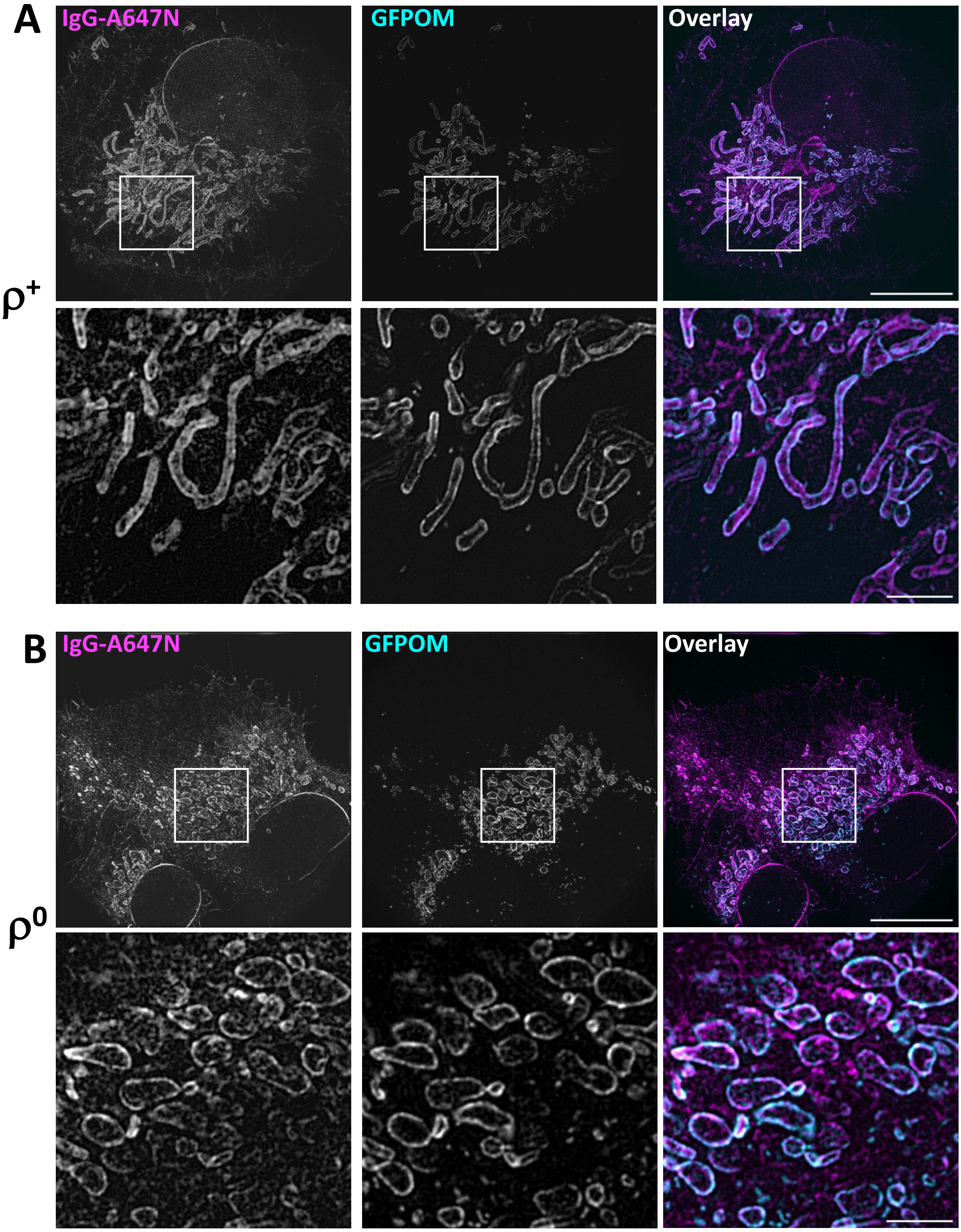


**Supplementary Figure S4: U-ExM of cells labelled with IgG-A647N unveils alterations of mitochondrial ultrastructure in $\rho^0$-cells devoid of mitochondrial DNA.** Stably transfected 143B cells expressing EGFP targeted to the outer membrane (GFPOM) were fixed, permeabilized, decorated with IgG-A647N, polymerized into a U-ExM gel and homogenized by partial proteolysis. **A**: Wild-type 143B $\rho^+$ cells: GFPOM is restricted to the boundary membrane. IgG-A647N labels mitochondrial inner and boundary membranes, **B**: 143B $\rho^0$ cells devoid of mitochondrial DNA: IgG-A647N appears restricted to GFPOM-positive boundary membranes. Bar 10 µm (main) and 2 µm (inset).
